# ASXL3 controls cortical neuron fate specification through extrinsic self-renewal pathways

**DOI:** 10.1101/2021.07.20.452995

**Authors:** BT McGrath, P Wu, S Salvi, N Girgla, X Chen, J Zhu, R KC, YC Tsan, A Moccia, A Srivastava, X Zhou, SL Bielas

## Abstract

During corticogenesis, transcription plasticity is fundamental to the restriction of neural progenitor cell (NPC) multipotency and production of cortical neuron heterogeneity. Human and mouse genetic studies have highlighted the role of Polycomb transcriptional regulation in this process. *ASXL3*, which encodes a component of the Polycomb repressive deubiquitination (PR-DUB) complex, has been identified as a high confidence autism spectrum disorder (ASD) risk gene. Genetic inactivation of *Asxl3,* in a mouse model that carries a clinically relevant *ASXL3* frameshift (*Asxl3^fs^)* variant, disrupts lateral expansion of NPCs and delays cortical neuron differentiation. Single-cell RNA sequencing analysis implicates Notch signaling, which alters the composition of excitatory neurons and fidelity of cortical layer deposition. Our data provides a new link between extrinsic signaling cues and intrinsic epigenetic regulation that together control the timing of cell fate programs. Furthermore, transcriptomic analysis revealed dysregulation of other known ASD risk genes indicating that a convergent developmental pathway is affected. Collectively our work provides important insights about developmental mechanisms that contribute to ASD neuropathology.

## INTRODUCTION

The cellular complexity of the neocortex is precisely orchestrated during development by the sequential generation of cortical neurons from a common pool of neural progenitor cells (NPCs) [1-8]. Progressive restriction of NPC multipotency, by highly defined gene expression patterns, determines the composition of excitatory cortical neurons distributed across the six-layers of the cerebral cortex. Chromatin organization and dynamic epigenomic regulation of histone modifications are required for the transcriptional plasticity that affords progressive restriction of NPC multipotency, while instilling mature cortical neuron fate on immature postmitotic neurons [9]. Genes that encode components of chromatin modification complexes comprise a prominent genetic etiology of developmental syndromes and autism spectrum disorders (ASD), further implicating an important role for chromatin biology in corticogenesis and neuron fate determination [10-12].

Polycomb group (PcG) complexes are a source of chromatin modifications that play an essential role in repressing gene expression to control development [13]. Much of our understanding of Polycomb transcriptional repression comes from characterization of Polycomb repressive complex 1 (PRC1) and Polycomb repressive complex 2 (PRC2). PRC1 monoubiquitylates histone H2A at lysine 119 (H2AUb1), and PRC2 mono-, di-, and tri-methylates histone H3 at lysine 27 (H3K27me1/2/3). Polycomb repressive deubiquitination complex (PR-DUB) is responsible for H2AUb1 deubiquitination, but is less well studied than PRC1 and PRC2 [14]. PRC1 and PRC2 loss-of-function studies support a role for PcG epigenetic repression in progressive restriction of NPC multipotency required for the full spectrum of mature cortical neuron production during corticogenesis [15, 16]. Human neurogenetic evidence suggest an important role for PR-DUB in brain development, with genes that encode PR-DUB complex components identified as the genetic basis of developmental disorders [17-21].

*De novo* dominant pathogenic variants in the PcG protein *ASXL3 (Additional sex comb-like 3)* were identified as the genetic basis of Bainbridge-Ropers syndrome and ASD [19]. H2AUb1 dysregulation is a key feature of ASXL3 molecular pathology and re-distribution of genome wide H2AUb1 is accompanied by altered transcriptional profiles [17]. ASXL3 expression is highest in the brain during the first 24 weeks of gestation (Brainspan, Allen Brain Institute), a period of cortical development characterized by rapid NPC proliferation, birth of postmitotic neurons, and specification of cortical neuron fates [22, 23]. The impact of ASXL3-dependent H2AUb1 deubiquitination on neural development has not been previously described.

Here we investigate the role of ASXL3 in modulating transcriptional programs that contribute to the neuronal diversity of the cortex. We created an *Asxl3* mouse model carrying a clinically relevant frameshift allele (*Asxl3^fs/fs^*). We detect elevated levels of H2AUb1, differential gene expression, and a cortical neuron fate defect that opposes the cortical phenotype described for conditional deletion of Ring1b, the catalytic component of PRC1 [15]. Characterization of cellular composition by single-cell RNA sequencing (scRNA-seq) at three developmental stages depicted an increase in *Asxl3^fs/fs^* neural progenitors, which was validated by immunohistochemistry and attributed to expansion of ventricular radial glia consistent with altered lateral expansion. Transcriptomic analysis implicates perturbations to extrinsic signaling pathways that disrupt the balance of NPC proliferation and differentiation. Transcriptional pathways are enriched with differentially expressed genes (DEGs) that converge on chromatin biology and proliferation at embryonic day (E) 13.5 and E14.5, and evolved to implicate synaptic regulation at birth. Across cortical development, ASD risk genes are represented in DEGs, implicating a convergence of pathogenic mechanisms among chromatin etiologies of ASD.

## RESULTS

### *Asxl3^fs/fs^* mice display multiple defects of nervous system development

To investigate the role of *Asxl3* in brain development, we generated a clinically relevant *Asxl3 c.990_992delCA* frameshift allele (fs) in mouse exon 12 using CRISPR/Cas9 genome-editing. *Asxl3* exhibits low, but persistent expression in developing mouse cortex starting as early as E10.5 [24] (Supplemental Fig. 1). *Asxl3^fs/fs^* mRNA is also detected in developing cortical tissue, but full length ASXL3 protein was not detected by western blot (Fig. 1A and Supplemental Fig. 1). Loss of ASXL3 is accompanied by a 3-fold increase of genome-wide H2AUb1 in the E13.5 cerebral cortex (Fig. 1B, C). Hetero- and homo-zygous *Asxl3* frameshift mice are born at Mendelian ratios. *Asxl3^+/fs^* mice are indistinguishable from *Asxl3^+/+^* at birth and breed normally. However, homozygous *Asxl3^fs/fs^* are perinatal lethal, with a majority of *Asxl3^fs/fs^* pups dying within the first week of life [25]. Multiple organ systems, including the heart, brain and skeleton are affected in *Asxl3^fs/fs^* mice. Hypoplastic heart ventricles is the predicted cause of perinatal death of *Asxl3^fs/fs^* animals [25]. *Asxl3^fs/fs^* mice exhibit a distinguished posture at birth with a curved back and drooping forelimbs (Fig. 1D). Alizarin red and alcian blue skeletal survey revealed a partially penetrant cervical rib in *Asxl3^fs/fs^* mice consistent with a homeotic transformation attributed to ectopic *Hox* gene expression, but ruled out kyphosis due to axial skeletal abnormalities [25]. Such posture has been observed in transgenic mouse models that disrupt neuromuscular axis elements, including upper motor neurons, corticospinal tracts, lower motor neurons and/or the neuromuscular junction [26].

**Fig. 1.**
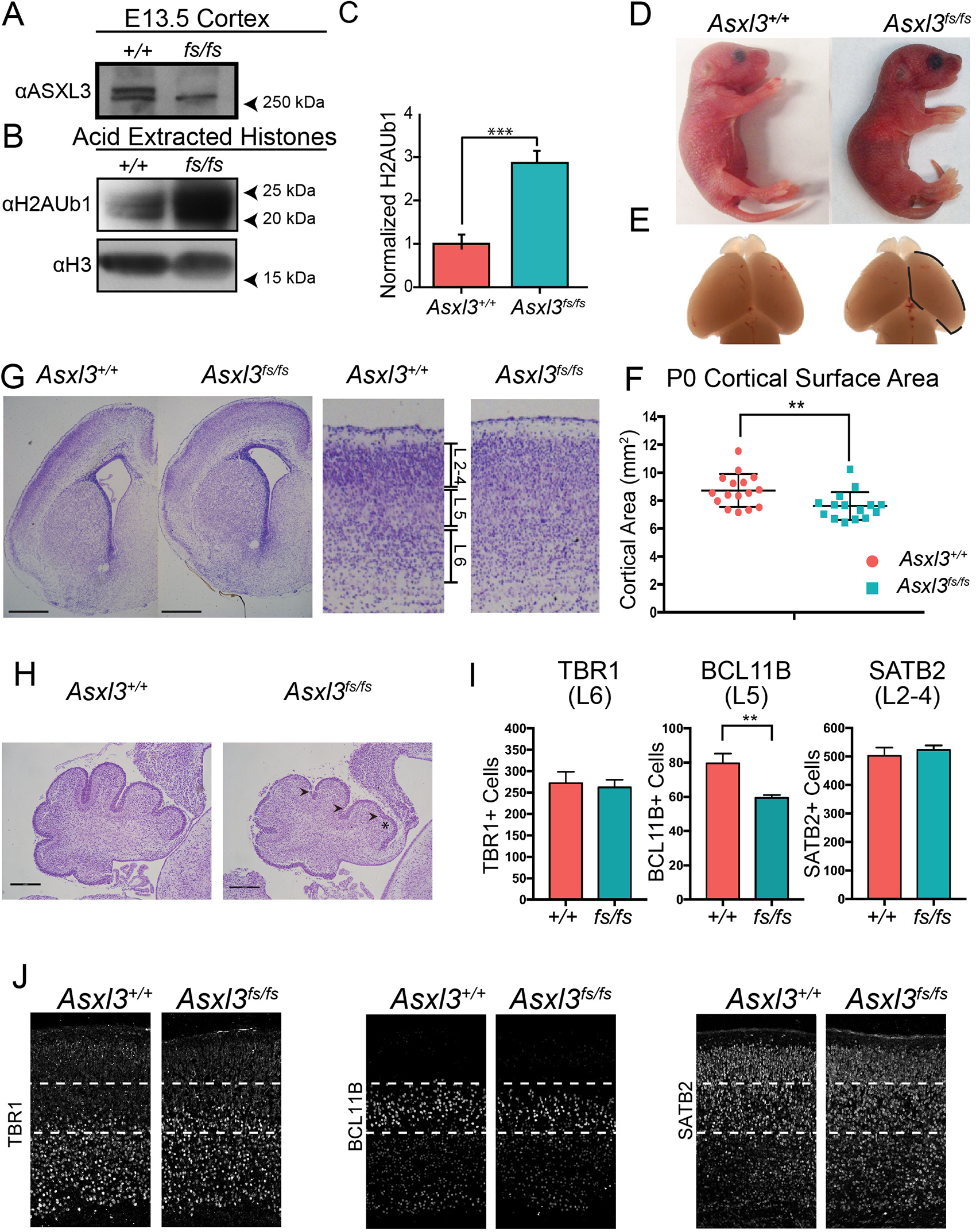
Loss of *Asxl3* disrupts cerebral cortex and cerebellum morphology A, Lysates from E13.5 *Asxl3^+/+^* and *Asxl3^fs/fs^* cortices immunoblotted for full length ASXL3. **B,** Acid extracted histones from E13.5 *Asxl3^+/+^* and *Asxl3^fs/fs^* E13.5 cortices immunoblotted for H2AUb1 and histone H3. **C,** Quantification of H2AUb1 levels relative to H3 in *Asxl3^+/+^* (*n*=3) and *Asxl3^fs/fs^* (*n*=3) E13.5 cortices.\*\*\**p*<0.001 using two-tailed unpaired Student’s *t* test. Values are displayed as mean ± SEM. **D,** Appearance of curved back and drooping forelimbs in P0.5 *Asxl3^fs/fs^* mice. **E,** Representative images and **F,** quantification of P0.5 cortical surface area, marked by a black dashed line, for *Asxl3^+/+^* (*n*=16) and *Asxl3^fs/fs^* (*n*=15) mice. *p*=0.009 using two-tailed unpaired Student’s *t* test. Values are displayed as mean ± SEM. **G,** Cresyl violet staining of *Asxl3^+/+^* vs *Asxl3^fs/fs^* P0.5 cerebral cortex. Whole hemisphere (left) and magnification of the cortex (right). Scale bar, 500 µm. **H,** Sagittal P2 cerebellar sections stained with cresyl violet. *Asxl3^fs/fs^* cerebellums have shallower fissures (arrowheads) and smaller lobules (asterisk). Scale bar, 500 µm. **I,** Quantification of the number of neurons expressing TBR1, BCL11B, or SATB2 in *Asxl3^+/+^* (*n*=4, *n*=8, *n*=3) and *Asx3^fs/fs^* (*n*=4, *n*=8, *n*=3) cortices. *p*=0.76 (TBR1), *p*=0.004 (BCL11B), *p*=0.57 (SATB2) using two-tailed unpaired Student’s *t* test. **J,** Immunohistochemistry of *Asxl3^+/+^* versus *Asx3^fs/fs^* P0.5 coronal cortical sections using layer-specific markers TBR1 (layer 6), BCL11B (layer 5), and SATB2 (layer 2-4).

Genetic disruption of individual PcG components give rise to a full spectrum of cortical size and cytoarchitectural changes [27]. *Asxl3^fs/fs^* cortical surface area and telencephalic hemisphere length were evaluated relative to control littermates. At P0, *Asxl3^fs/fs^* cortical surface area is 87% of *Asxl3^+/+^* littermates, while hemispheric length is not significantly shorter (Fig. 1E, F and Supplemental Fig. 1). Gross structural defects in cortical anatomy or commissure formation were not observed in a series of matched coronal and sagittal sections of *Asxl3^fs/fs^* and *Asxl3^+/+^* P0.5 paraffin fixed brains (Fig. 1G and Supplemental Fig. 1). Sagittal sections through the cerebellar hemispheres reveal shallower fissures and smaller lobules in *Asxl3^fs/fs^* versus *Asxl3^+/+^* P2 littermates consistent with a delay in cerebellar development (Fig. 1H). *Asxl3^fs/fs^* cortices lack distinct boundaries between cortical layers in cresyl violet stained sections (Fig. 1G). To explore this phenotype, cortical layers (L) were evaluated by immunohistochemistry (IHC) using markers of mature neuron subtypes, including TBR1, BCL11B, and SATB2 (Fig. 1J and Supplemental Fig. 2). A >20% reduction of BCL11B labeled L5 neurons was quantified in the P0.5 cortex of *Asxl3^fs/fs^* mice relative to *Asxl3^+/+^* controls (Fig. 1I). This phenotype does not correlate to a corresponding change in the number of L6 TBR1 and/or L2-4 SATB2 cortical neurons.

### Lateral expansion generates increased NPCs in *Asxl3^fs/fs^* cortex

During cortical development, NPCs sequentially give rise to distinct cortical neuron subtypes. To characterize the developmental source of P0.5 cortical neuron composition changes, scRNA-seq using Seq-Well was performed on cortices from E13.5 and E14.5 littermates. NPCs at these developmental time points give rise to L5 neurons. In total, 17 cortices were processed across these two timepoints (Supplemental Fig. 3). Following quality control steps, data from *Asxl3^+/+^* and *Asxl3^fs/fs^* samples at E13.5 and E14.5 were integrated and analyzed using Seurat [28]. Standard quality control features for brain based single cell transcriptomics and Seq-Well studies were used (Supplemental Fig. 3) [29-31] Clustering was not driven by experimental batch or individual samples (Supplemental Fig. 4) indicating consistent transcriptional signatures for individual cell types between animals. After performing dimensionality reduction and unsupervised clustering with uniform manifold approximation and projection (UMAP) we identified unique cell populations for each developmental timepoint (Supplemental Fig. 4). 46,920 E13.5 and 24,963 E14.5 cortical cells were retained that span twelve and fifteen clusters respectively (Supplemental Fig. 3 and Fig. 4). *Asxl3^+/+^* and *Asxl3^fs/fs^* cells were represented in all clusters (Supplemental Fig. 4). Cluster cell-type identities were manually annotated based on cluster-specific molecular markers as defined by previous studies [29, 30, 32]. We confirmed our annotation by aligning our *Asxl3^+/+^* E14.5 dataset with published E14.5 mouse brain scRNA-seq data [29]. Cluster assignments between the two datasets were closely matched, with little representation of cell types outside our dissection perimeter (Supplemental Fig. 5). This highlights the reproducibility of scRNA-seq between studies and the robustness of our data to characterize developmental defects.

Following cell-type annotation, we compiled transcriptomic data from clusters which contribute to excitatory lineages. Unsupervised clustering of data from this subset of cells was performed and clusters annotated (Fig. 2). At E13.5, eleven clusters were identified corresponding to known excitatory lineage cell types, including seven radial glia cell (RGC), a differentiating radial glia cell (dRGC), an intermediate progenitor (IP), a migrating neuron, and two L5-6 excitatory neuron (DL neuron) clusters (Fig. 2B). At E14.5, ten clusters were defined that included four RGC, a dRGC, two IP, a migrating neuron, and three DL neuron populations (Fig. 2E). Consistent with scRNA-seq data from developing human cortex, the clustering resembled the predicted developmental trajectory with RGCs transitioning to IPs then migrating neurons and finally DL neurons (Fig. 2E) [30]. We next examined the distribution of canonical marker genes that define cells at different stages of neurogenesis (Fig. 2G, H and Supplemental Fig. 6). The numerous RGC populations (Clusters E13-0, 13-1, 13-2, 13-3, 13-4, 13-5, 13-6, 14-0, 14-1, 14-2, 14-3) express a combination of cell cycle genes like *MKi67*, *Aspm*, *Lig1* and RGC transcription factors like *Pax6* and *Sox2*. We observed that dRGC (Clusters E13-7, 14-4) co-express markers of the other major cell types (RGC, IP, migrating neuron, DL neuron) at a reduced level. Similar transitory cells have been described in transcriptomic studies of developing mouse and human cortex [30, 33, 34]. IP clusters (Clusters E13-8,14-5,14-6) are marked by *Eomes*. Migrating neurons express a combination of general neuronal markers *Neurod2*, *Neurod6*, and axon guidance genes *Rnd2*, *Cntn2*, and *Robo2*. Among L5-6 excitatory neuron clusters, expression of transcription factors that demark deep and upper layer excitatory neurons was noted, such as *Sox5*, *Tbr1*, *Bcl11b*, *Bcl11a*, *Fezf2*, *Satb2*, and *Pou3f1*.

**Fig. 2.**
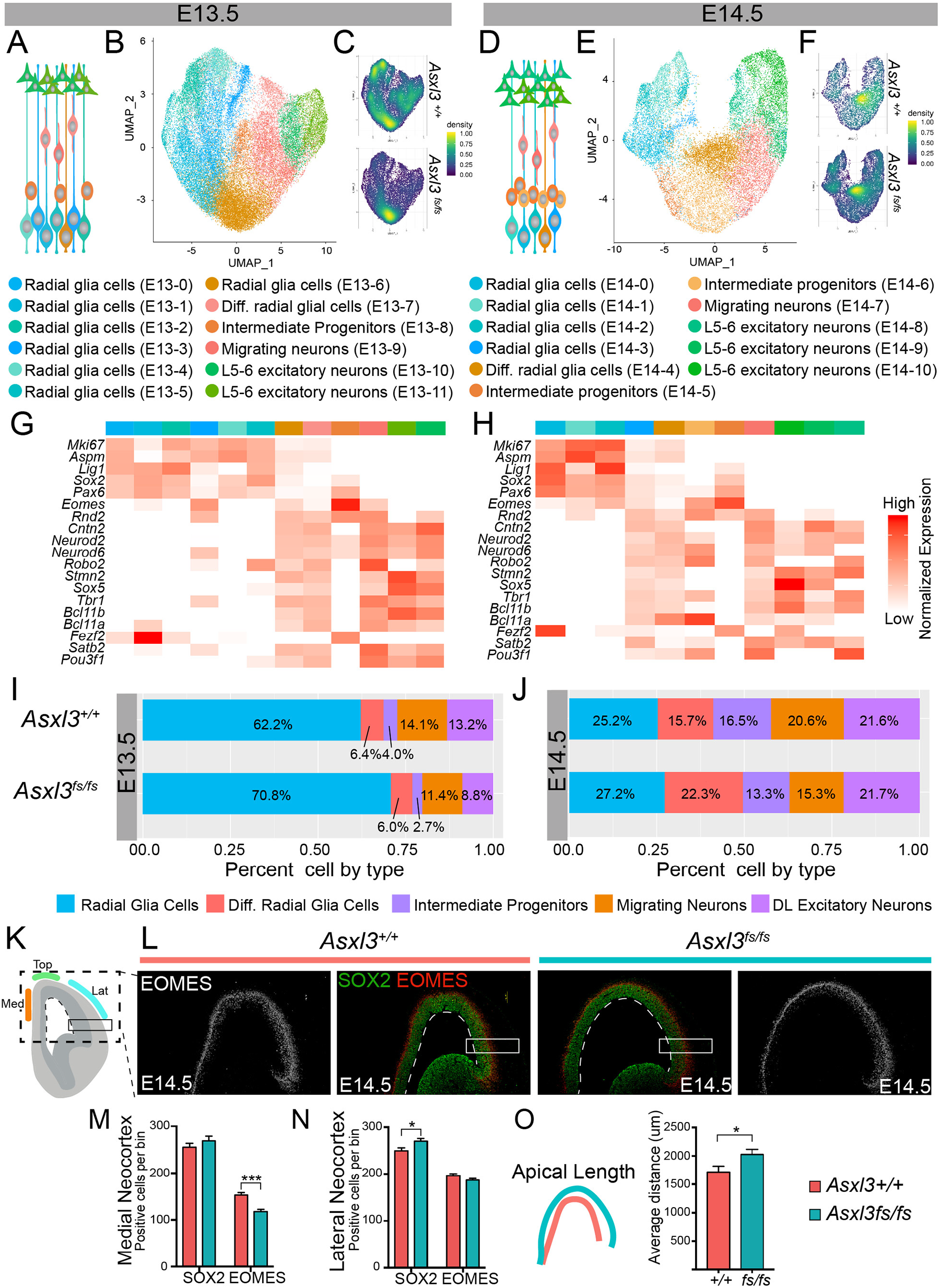
Neural progenitor cells expanded in *Asxl3^fs/fs^* cortices A, Depiction of E13.5 cortical excitatory neurons. **B,** Unsupervised clustering of E13.5 excitatory neurons collected from *Asxl3^+/+^* (26,780 cells; *n*=5) and *Asx3^fs/fs^* (25,144 cells; *n*=3) mice color coded by cell type and represented on a UMAP. **C,** Distribution of *Asxl3^+/+^* (top) and *Asx3^fs/fs^* (bottom) cells across populations in the E13.5 UMAP. **D,** Depiction of E14.5 cortical excitatory neurons. **E,** Unsupervised clustering of E14.5 excitatory neurons collected from *Asxl3^+/+^* (8,261 cells; *n*=5) and *Asx3^fs/fs^* (22,075 cells; *n*=7) mice color coded by cell type and represented on a UMAP. **F,** Distribution of *Asxl3^+/+^* (top) and *Asx3^fs/fs^* (bottom) cells across populations in the E14.5 UMAP. The UMAPs in C and F are colored by density with yellow indicating a high density and blue a low density. Heatmap showing normalized expression of key marker genes expressed in **G**, E13.5 and **H**, E14.5, cortices. Comparison of the proportion of major cell types detected in the *Asxl3^+/+^* and *Asx3^fs/fs^* datasets at **I**, E13.5, and **J**, E14.5. Major cell types include radial glia cells (RGC E13.5 *p=*6.3e^-79^, E14.5 *p*=0.02), differentiating radial glia cells (dRGC E13.5 *p=*0.1, E14.5 *p*=1.7e^-72^), intermediate progenitors (IP E13.5 *p=*8.1e^-12^, E14.5 *p*=2.8e^-09^), migrating neurons (E13.5 *p=*4.2e^-16^, E14.5 *p*=2.6e^-20^), and deep layer excitatory neurons (E13.5 *p=*3.6e^-46^, E14.5 *p*=0.82). P values were generated using a two proportion Z test. **K,** Schematic illustrating quantifications collected from immunostained coronal sections of the cortex. SOX2 and EOMES positive cells were counted in bins (black boxes) throughout the medial, top, and lateral neocortex. Apical length (white dashed line) was measured throughout serial sections. **L,** Immunohistochemical staining of *Asxl3^+/+^* (left) and *Asx3^fs/fs^* (right) E14.5 coronal cortical sections with SOX2 (Green) and EOMES (grey, red). White boxes denote regions of quantification. White dashed line marks the apical length. Quantification of the number of cells expressing SOX2 or EOMES in the **M,** medial, or **N,** lateral neocortex in *Asxl3^+/+^* (*n*=6) and *Asx3^fs/fs^* (*n*=6) mice. M, *p*=0.3 (SOX2), *p*=1.3×10^-6^ (EOMES). N, *p*=0.02 (SOX2), *p*=0.11 (EOMES) using two-tailed unpaired Student’s *t* test. **O,** Quantification of the apical length of E14.5 *Asxl3^+/+^* (*n*=8) and *Asx3^fs/fs^* (*n*=8) littermate cortices. *p*=0.038 using two-tailed unpaired Student’s *t* test. In M-N values are displayed as mean ± SEM.

While *Asxl3^fs/fs^* and *Asxl3^+/+^* cells are distributed across all clusters at E13.5, the density of *Asxl3^fs/fs^* cells are significantly enriched in the region of UMAP space that overlaps with cluster E13-6 (Fig. 2C and Supplemental Fig. 4C). Cells in E13-6 have the transcriptional characteristics of a transition cell type, uniformly expressing markers of RGC and other major cell types at low levels (Fig. 2G). The position of cluster E13-6 in UMAP space suggest the cluster is composed of RGC cells differentiating to IPs and/or immature postmitotic migrating neurons. These findings are consistent with a model where progenitors exhibit a permissive transcriptional status that supports gradual differentiation to excitatory linage fates, independent of a direct or indirect path through IPs [35, 36]. At E14.5, all clusters were populated by *Asxl3^fs/fs^* and *Asxl3^+/+^* cells, without genotype-dependent enrichment in one or more cell-specific clusters (Fig. 2E and Supplemental Fig. 4B).

To assess changes in *Asxl3^fs/fs^* cortical cell composition, the overall proportions of developmental cell classes were calculated at E13.5 and E14.5. For this analysis cortical cells were grouped into five classes, namely RGC, dRGC, IP, Migrating neurons, and DL neurons. When compared to *Asxl3^+/+^*, RGCs are significantly increased at E13.5 in *Asxl3^fs/fs^* cortex, with a corresponding decrease in more mature cell types (Fig. 2I and Supplemental Fig. 4F). A day later at E14.5, the E13.5 increased RGCs has transitioned to include more dRGCs and fewer IPs and migrating neurons in *Asxl3^fs/fs^* cortex (Fig. 2J and Supplemental Fig. 4E). This change implicates dysregulation of RGC proliferation or differentiation dynamics in *Asxl3^fs/fs^* developing cortex.

To characterize the developmental mechanism that accounts for *Asxl3^fs/fs^* cellular composition changes, we immunostained E14.5 cortical tissue for SOX2 and EOMES, to evaluate RGCs and IPs (Fig. 2K, I). Altered proliferation dynamics can alter NPC niche size by modulating ventricular (VZ)/subventricular (SVZ) zone thickness (radial expansion) or ventricular zone length (lateral expansion). Both radial and lateral VZ changes were quantified. To evaluate VZ/SVZ thickness, the number of SOX2+ RGCs and EOMES+ IPs in a single bin that spans the cortical thickness was quantified at medial, medial/lateral and lateral regions of the neocortex (Fig. 2K). A small but significant radial expansion of SOX2+ RGCs was detected in the lateral neocortex and a decrease in EOMES+ IPs was identified in the medial and medial/lateral neocortex (Fig. 2M, N and Supplemental Fig. 7). These changes would not entirely account for *Asxl3^fs/fs^* progenitor increase detected by scRNA-seq. Lateral expansion was assessed by measuring the length of the lateral ventricular/VZ boundary that corresponds to the apical border of the VZ. The length of the *Asxl3^fs/fs^* VZ was significantly longer along all rostral to caudal sections measured (Fig. 2O and Supplemental Fig. 7). Therefore, *Asxl3^fs/fs^* cortical cell composition changes detected by scRNA-seq can largely be attributed to lateral expansion of VZ RGCs. Similar phenotypes have been observed in mouse models that disrupt signaling pathways critical for balancing NPC maintenance and differentiation [37-39].

### Deep layer cortical neuron differentiation delayed in *Asxl3^fs/fs^* cortex

Corticogenesis relies on an exquisite balance between RGC expansion and neuronal differentiation that generates the full complement of mature cortical neuron subtypes. The embryonic increase in *Asxl3^fs/fs^* progenitors suggests the balance of this stereotypic process is disrupted. Pseudotemporal analysis was performed using Monocle 3 to reconstruct and evaluate the differentiation process of E13.5 and E14.5 corticogenesis (Fig. 3A-F) [40]. At both time points, pseudotime ordering of cell types mirrored *in vivo* corticogenesis. RGCs represent the earliest pseudotime stages of differentiation followed by dRGCs, IP, Migrating neurons, and DL excitatory neurons (Fig. 3A, D and Supplemental Fig. 8). We then constructed individual trajectories for *Asxl3^+/+^* and *Asxl3^fs/fs^* samples to compare genotypic pseudotime differentiation outcomes (Fig. 3B, E). Again, at E13.5 an *Asxl3^fs/fs^* increase in RGCs and corresponding decrease in more mature cell types was noted, based on a quantitative change in the amplitude, or cell density, of overlapping pseudotime peaks. This difference in pseudotime landscape is consistent with RGC multipotency persistence at the expense of neurogenic divisions. A genotypic dependent change in the position of peaks across pseudotime, consistent with a change in the timing of differentiation was not observed at E13.5 (Fig. 3B-C). E14.5 analysis revealed *Asxl3^fs/fs^* changes in amplitude and distribution of cell type-specific peaks. An increase in progenitors persists, but is now distributed across RGC, dRGC and IP cell types (Fig. 3E, F). However, progenitor persistence now correlates with a novel *Asxl3^fs/fs^* peak detected between pseudotime differentiation position 20-30. The peak exhibits unique spacing relative to *Asxl3^+/+^* peaks and is composed of migrating neurons and DL excitatory neurons, consistent with delayed differentiation of a subset of *Asxl3^fs/fs^* deep layer cortical neurons (Fig. 3E, F). Collectively, the analysis supports a model where persistence of RGC self-renewing proliferation at E13.5 manifests in delayed differentiation of a population of excitatory neurons evident at E14.5.

**Fig. 3.**
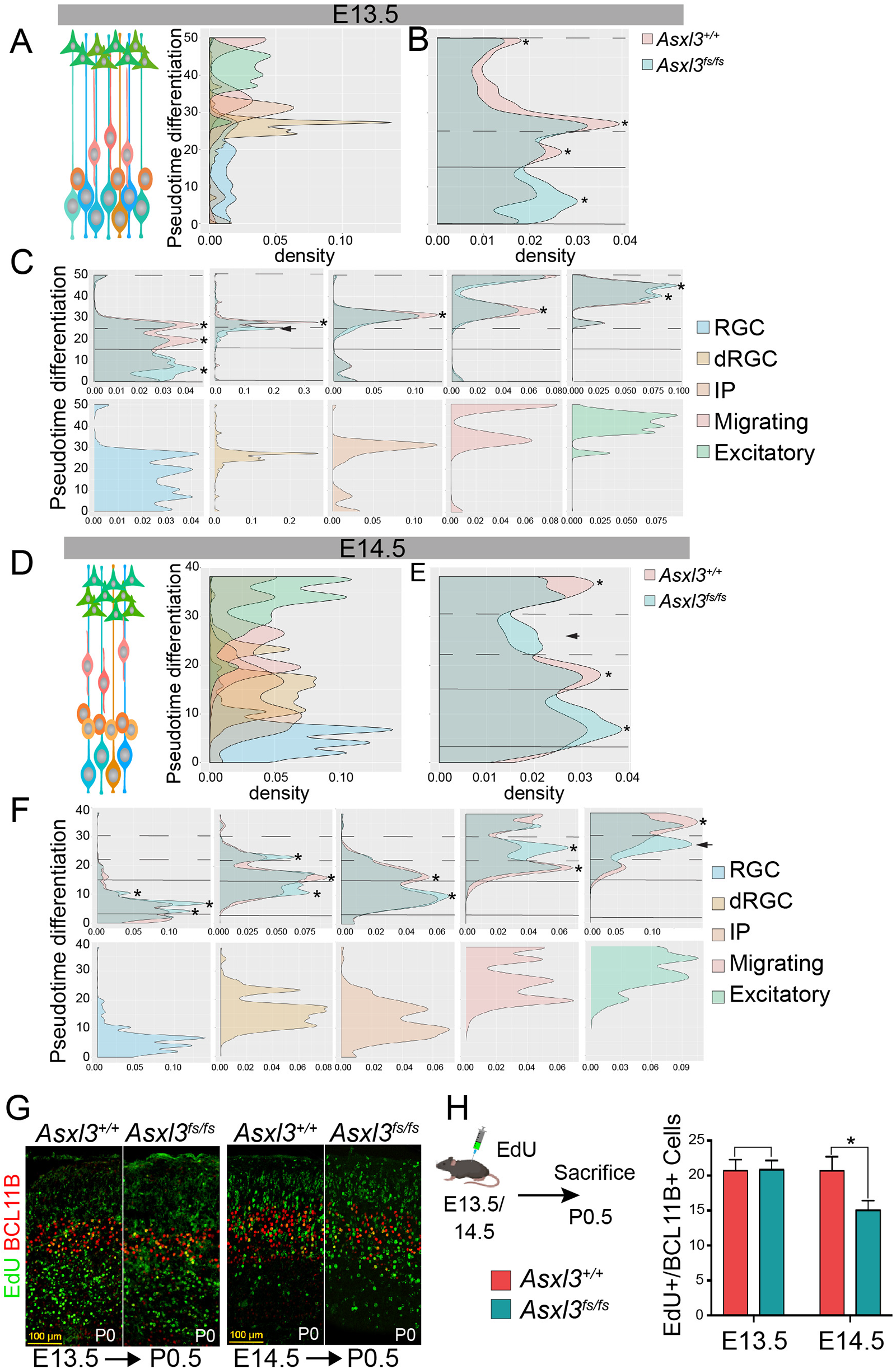
*Asxl3^fs/fs^* cortices show altered timing of neuronal differentiation A, Histograms displaying number of E13.5 cells ordered along pseudotime colored by cell type. **B,** E13.5 pseudotime histogram colored by genotype (*Asxl3^+/+^*, red; *Asxl3^fs/fs^*, blue). **C,** Individual E13.5 pseudotime histograms of RGC, dRGC, IP, Migrating neurons, and DL excitatory neurons. **D,** Histograms of the number of E14.5 cells ordered along pseudotime colored by cell type. **E,** E14.5 pseudotime histogram colored by genotype (*Asxl3^+/+^*, red; *Asxl3^fs/fs^*, blue). **F,** Individual E14.5 pseudotime histograms of RGC, dRGC, IP, Migrating neurons, and DL excitatory neurons. An asterisk indicates peaks with changes in amplitude and arrowheads indicate peaks with a shift in pseudotime. Birthdating analysis was performed at E13.5 and E14.5. EdU was injected at either E13.5 or E14.5 and then labeled brains were collected at P0.5. **G,** Representative P0.5 *Asxl3^+/+^* and *Asxl3^fs/fs^* cortical sections after EdU detection and immunostained with BCL11B. The left panel shows sections from animals injected with EdU at E13.5 and the right shows E14.5. **H,** Quantification of EdU+/BCL11B+ cells in E13.5 injected cortices (*p*=0.947) and E14.5 (*p*=0.018) using two-tailed unpaired Student’s *t* test. values are displayed as mean ± SEM.

To assess the relationship of progenitor expansion on neuron differentiation, E13.5 or E14.5 time-pregnant dames were administered EdU and sacrificed at P0.5. The majority of EdU labeled cells at E13.5 were distributed across deep cortical layers, with a comparable distribution in both genotypes. No change in the percentage of EdU labeled or EdU/BCL11B double-labeled cells at E13.5 was quantified (Fig. 3G, H). At E14.5, the distribution of EdU labeled cells included upper cortical layers in *Asxl3^+/+^* littermates, with few cells labeled below BCL11B L5 neurons. *Asxl3^fs/fs^* EdU labeled cells exhibit greater distribution across cortical layers, including deep layers and a reduction in the percentage of EdU/BCL11B double-labeled cells was quantified in *Asxl3^fs/fs^* cortex, consistent with a delay in cortical neuron differentiation.

### Altered extrinsic signaling disrupted in *Asxl3^fs/fs^* corticogenesis

To uncover the cell and molecular biology impacted by *Asxl3^fs/fs^* transcriptional changes and that may account for the corticogenesis defects, Gene Ontology (GO) analysis of E13.5 and E14.5 clusters was performed. Corresponding E13.5 and E14.5 *Asxl3^fs/fs^* DEGs were enriched in GO terms implicating extrinsic signaling pathways, including Notch, Wnt, and Hippo (Fig. 4A). Enriched DEGs of note include *Hes1*, *Hes5*, *Notch1, Notch2, Dll1, Wnt5a, Wnt7b, Fzd1, Fzd4, Yap1, Tead1, Tead2*. Activities of these extrinsic signaling pathways have been demonstrated to act on NPCs to direct the balance between NPC proliferation and differentiation [37, 39, 41, 42]. Mouse models of constitutive overexpression or persistent activity have demonstrated cross talk between these signaling pathways during cortical development [37, 41].

DEGs in the Notch signaling pathway were used to investigate how coordination of temporally defined cell-specific transcription cascades are dysregulated to contribute to *Asxl3^fs/fs^* corticogenesis defects. Notch function is mediated through cell-cell interactions of neighboring cells (Fig. 4B). Expression of a transmembrane ligand (delta or jagged) in one cell, binds a Notch receptor on an opposing cell. A series of cleavage events allows the Notch intracellular domain (NICD) to translocate into the nucleus. Intranuclear NICD initiates transcription of bHLH transcription factors *Hes1/5* that inhibit neuronal differentiation by repressing expression of proneural bHLH *Neurog2*. In mice, overexpression of N1ICD, *Hes1* or *Hes5* in the developing cortex induces lateral expansion of NPCs [37, 38, 43]. N1ICD ChIP-seq and transcriptomic analysis of the N1ICD overexpression mouse cortex identified direct and downstream targets of N1ICD [37]. These targets were intersected with E13.5 and E14.5 individual cluster transcriptional changes which revealed enrichment for direct and downstream N1ICD targets amongst the *Asxl3^fs/fs^* DEGs (Fig. 4C, D).

Significant *Hes1* and *Hes5* differential overexpression in E13.5 RGC clusters and phenocopy between *Asxl3^fs/fs^* and *Hes1* and *Hes5* overexpression mouse models implicate disruption of Notch transcription factor coordination as a molecular driver of early *Asxl3^fs/fs^* cortical defects (Fig. 4C, D). Supporting this observation, Notch DEGs that regulate *Hes1* and *Hes5* expression were detected across E13.5 clusters, while few downstream targets of HES1/5 were among the DEGs identified (Fig. 4E). In line with disruption of coordinated Notch signaling, at E14.5 fewer regulators of *Hes* expression were differentially expressed in RGCs, *Hes1/5* expression was largely normalized, while an increase in downstream targets of *Hes1/5* were enriched among the DEGs (Fig. 4F). If we integrate the data from these two time points, it implicates a molecular mechanism where upregulation of *Hes* genes at E13.5 favors retention of multipotent RGC fate through a mechanism of lateral expansion. By E14.5 downstream targets of HES1/5 downregulate, such as *Neurog2* which participates in a negative feedback loop with the *Hes* genes in Notch signaling (Fig. 4B). These results would support a shift in the balance towards self-renewing progenitor proliferation and away from neurogenesis or neuronal differentiation.

**Fig. 4.**
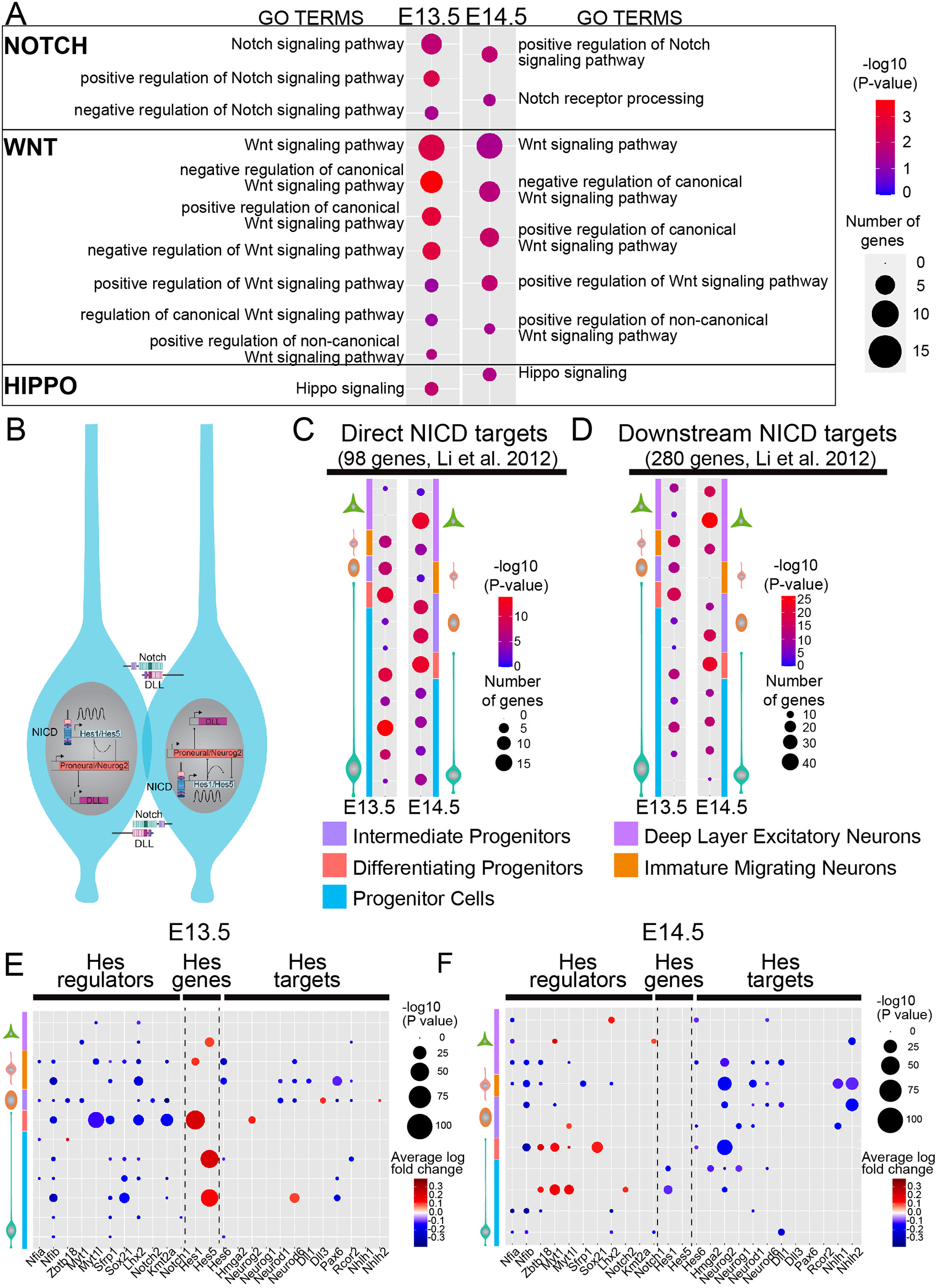
Extrinsic signaling pathways disrupted in *Asxl3^fs/fs^* developing cortex A, Gene ontology analysis showing statistically significant enrichment for dysregulated genes in E13.5 (left) and E14.5 (right) cortices. GO terms were enriched for Notch, Wnt, and Hippo genes. The size of each dot represents the gene number and the shading represents the -log_10_ *p*-value. **B,** Model of Notch signaling between neighboring RGC during neurogenesis to promote maintenance of progenitor multipotency. Binding of Notch ligands to the Notch receptors leads to a series of cleavage events and subsequent translocation of the NICD into the nucleus. Within the nucleus, the Notch intracellular domain (NICD) promotes the expression of *Hes1*/*Hes5.* In turn, HES1/HES5 repress *Neurog2* and other proneural genes that promote differentiation. Enrichment analysis for **C,** direct NICD and **D,** downstream NICD targets identified by Li et al. 2012 in our E13.5 and E14.5 differentially expressed genes. The size of each dot represents the number of genes and the shading represents the -log_10_ *p*-value. Each row represents enrichment of targets within DEGs from a single cluster. Results are grouped and colored by cell type. A Fisher’s exact test was used to determine if the gene lists were enriched. Cell type specific expression changes of Hes regulators, Hes genes and Hes targets based on scRNA-seq from **E,** E13.5 and **F,** E14.5 *Asxl3^fs/fs^* relative to *Asxl3^+/+^*. The size of each dot represents the -log_10_ *p*-value and the shading represents the average log fold change. Results are grouped and colored by cell type.

### Alterations in *Asxl3^fs/fs^* excitatory neuron composition

To characterize P0.5 cortical neuron composition by single cell transcriptomics, the P0.5 cortex was micro-dissected from brain stem, olfactory bulbs and ventral brain structures (Fig. 5A). scRNA-seq was performed on 23,511 cells from four *Asxl3^+/+^* and 38,672 cells from five *Asxl3^fs/fs^* cerebral cortices. Cells clustered into 21 unique cell populations and were annotated as described for embryonic timepoints (Supplemental Fig. 3). The cell populations identified were correlated to known major cell-types present in the P0.5 cortex, including subtypes of neural progenitors, intermediate progenitors, excitatory neurons and interneurons that are derived from ventral brain structures and tangentially migrate into the cortex (Supplemental Fig. 4). Comparable clusters were detected in both genotypes (Fig. 5C and Supplemental Fig. 4A). *Asxl3^+/+^* P0.5 data aligned with Loo et al. 2019 [29] P0 cortical scRNA-seq exhibited comparable clusters with minimal detection of cells from the striatum or ganglionic eminence, which were removed during dissection (Supplemental Fig. 5). This indicates the reproducibility of scRNA-seq between studies and the robustness of our data to characterize cortical neuron composition changes between *Asxl3^+/+^* and *Asxl3^fs/fs^* samples.

**Fig. 5.**
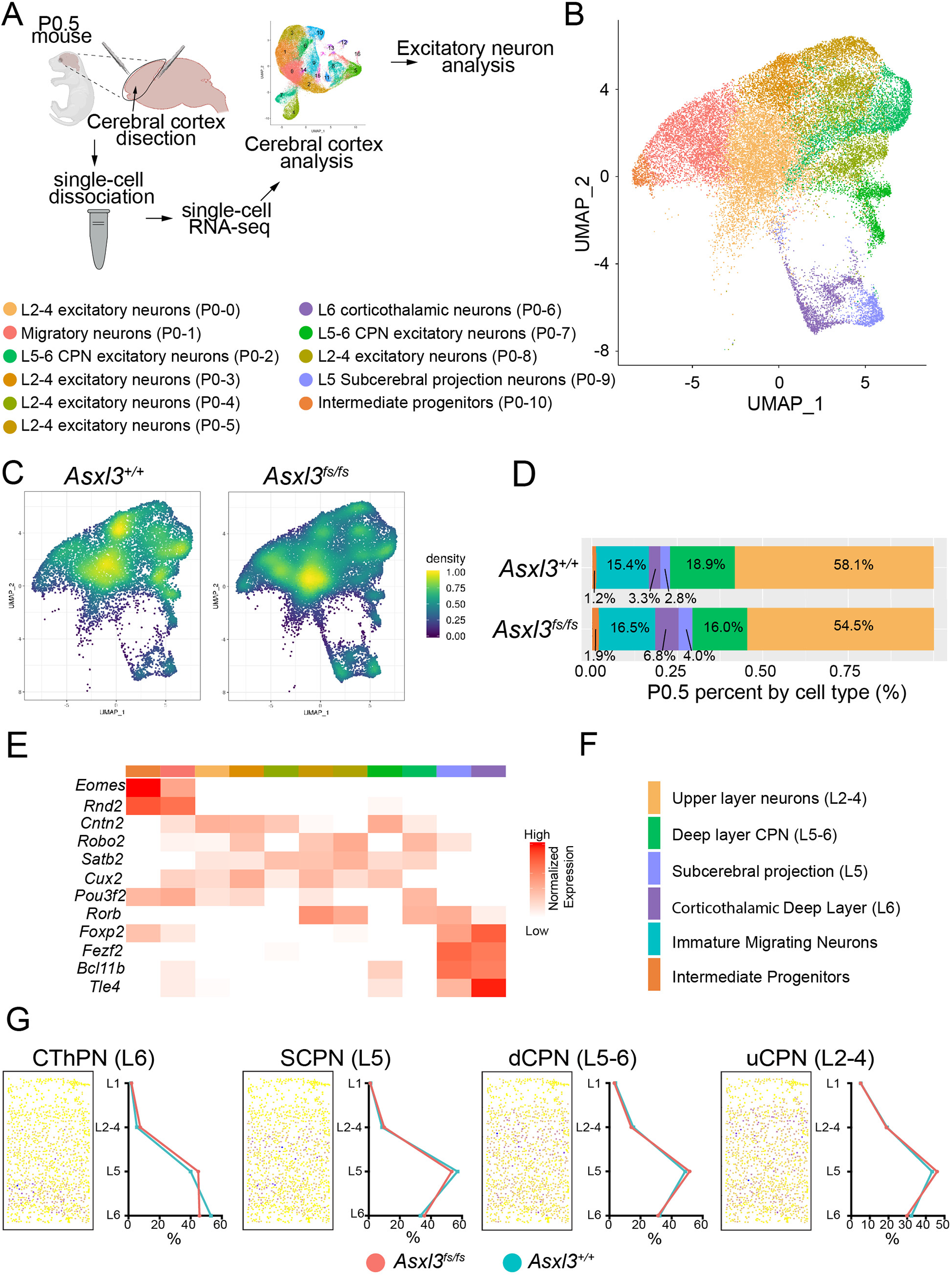
Excitatory cortical neuron composition altered in *Asxl3^fs/fs^* cortex A, Experimental procedure schematic depicting dissection of P0.5 cortex, single cell dissociation, scRNA-seq, and bioinformatic enrichment of excitatory neurons. **B,** Unsupervised clustering of P0.5 excitatory neurons collected from *Asxl3^+/+^* (31,798 cells; *n*=4) and *Asx3^fs/fs^* (44,384 cells; *n*=6) mice color coded by cluster and represented on a UMAP. Cell type annotations are located to the right. **C,** Distribution of *Asxl3^+/+^* (top) and *Asx3^fs/fs^* (bottom) cells across populations in the UMAP. The UMAP is colored by density with yellow indicating a high density and blue a low density. **D,** Comparison of the proportion of major cell types detected in the *Asxl3^+/+^* and *Asx3^fs/fs^* datasets. IP *p*=3.7e^-06^, Migrating *p*=0.01, CThPN *p*=5.2e^-37^, SCPN *p*=8.9e^-08^, dCPN *p*=7.7e^-11^, uCPN *p*=1.4e^-09^ using a two proportion Z test. **E,** Heatmap showing normalized expression of key marker genes expressed in P0.5 cortices. **F,** Major cell types and corresponding color code **G,** Left: mapping of our single cell gene expression data onto STARmap spatial gene expression data (Wang et al. 2018) using Tangram (Biancalani et al. 2020). Dots colored by the probability of our cell type mapping to the STARmap cell type (blue high probability, yellow low probability). Right: distribution of cells mapping to spatial gene expression data for CThPN, SCPN, dCPN, and uCPN.

To assess the excitatory neuron composition, transcriptomic data from excitatory neuron lineages was isolated from total cortical cell data (Fig. 5A). Comparable clusters were detected in both genotypes when clustered independently and in aggregate (Fig. 5C and data not shown). Eleven distinct clusters were annotated corresponding to IPs, Migrating neurons, and cortical pyramidal neurons from upper and deep layers, further classified as subcerebral projection neurons (SCPN), corticothalamic projection neurons (CThPN) and callosal projection neurons (CPN). Layer 5/6 deep layer CPNs (dCPN) were present in cluster P0-2 and P0-7. Layer 2-4 upper layer CPNs (uCPN) were in P0-0, P0-3, P0-4, P0-5, P0-8. Several marker genes for these CPN clusters have regional-specific expression including *Lhx2* (caudo-rostral), *Cbln2* (rostro-caudal), *Crym* (caudo-rostral), *Dkk3* (visual cortex), *Epha5* (somatosensory cortex), *Lmo4* (sensory areas), Bhlhb22 (Sensory areas) (Fig. 5F) [44]. Genes known to have subtype specific expression in CThPN and ScPN were enriched in clusters P0-6 and P0-9 respectfully (Fig. 5F and Supplemental Fig. 5).

The ratio of broad cell classes were calculated for *Asxl3^+/+^* and *Asxl3^fs/fs^* samples to determine the cortical composition of cells from excitatory lineages. Overall, deep layer cortical neurons classes exhibit the largest genotypic differences in cortical composition changes that diminished in upper layer neuron subtypes. In *Asxl3^fs/fs^* samples a two-fold increase in CThPNs coincides with a 30% increase in SCPNs and switches to a reduction in dCPNs (15% reduction) and uCPNs (6% reduction) relative to control (Fig. 5E). This finding is consistent with an early disruption to the timing of cortical neuron differentiation.

We evaluated the spatial distribution of *Asxl3^+/+^* and *Asxl3^fs/fs^* cells for individual combined excitatory clusters by employing Tangram (Fig. 5G and Supplemental Fig. 9) [45, 46]., Tangram maps cells identified through scRNA-seq by aligning the data with a reference spatial dataset, in our case the STARmap spatial dataset of the mouse visual cortex [46]. *Asxl3^+/+^* and *Asxl3^fs/fs^* P0.5 scRNAseq profiles were mapped based on 166 training genes present in both STARmap and our data. Of note, cell type distributions of excitatory neuron classes do not exhibit consistent layer-specific spatial patterns. CThPNs exhibit the greatest fidelity to the localized L6 spatial distribution in deep layers. Patterns of SCPNs, dCPNs and uCPNs distribution are less specific, with cells roughly distributed throughout upper and deep layers. These excitatory neuron classes were distinguished based on the percentage of cells mapped in L5-6 versus L2-4 (SCPN L6-34%. L5-56%. L2/4-9%; dCPN L6-32%, L5-50%, L2/4-15%; uCPN L6-31%, L5-44%, L2/4-19%). This distribution is consistent with growing literature implicating a greater contribution of shared transcriptional profiles along the continuum of excitatory neuron differentiation as fate is established [36, 47].

### ASXL3 disrupts differentiation of deep layer neurons

The P0.5 pseudotime differentiation trajectory of *Asxl3^+/+^* and *Asxl3^fs/fs^* cortical neuron subtypes was assessed as described for embryonic timepoints. The fundamental steps of cortical neuron differentiation are captured in this analysis, with undifferentiated IPs preceding Migrating neurons and terminally differentiated classes of excitatory neurons (Fig. 6A, B). The developmental relationship between cortical neuron subtypes along the pseudotime axis reflects the chronological acquisition of terminal fate. As such, early born L6 CThPN represent the most mature, or the terminus of differentiation and successive classes of later fated cortical neuron classes, SCPN, dCPN, uCPN, are correctly organized along the axis progressively closer to the immature cell types (Fig. 6C and Supplemental Fig. 9). This fidelity to the developmental process of corticogenesis allows differences in the individual genotypic trajectories to be identified. Most prominent among them are the differential *Asxl3^fs/^*^fs^ amplitude peak increase between pseudotime position 25-35 and the corresponding reduced *Asxl3^fs/fs^* amplitude peak between differentiation position 15-25 (Fig. 6A).

**Fig. 6.**
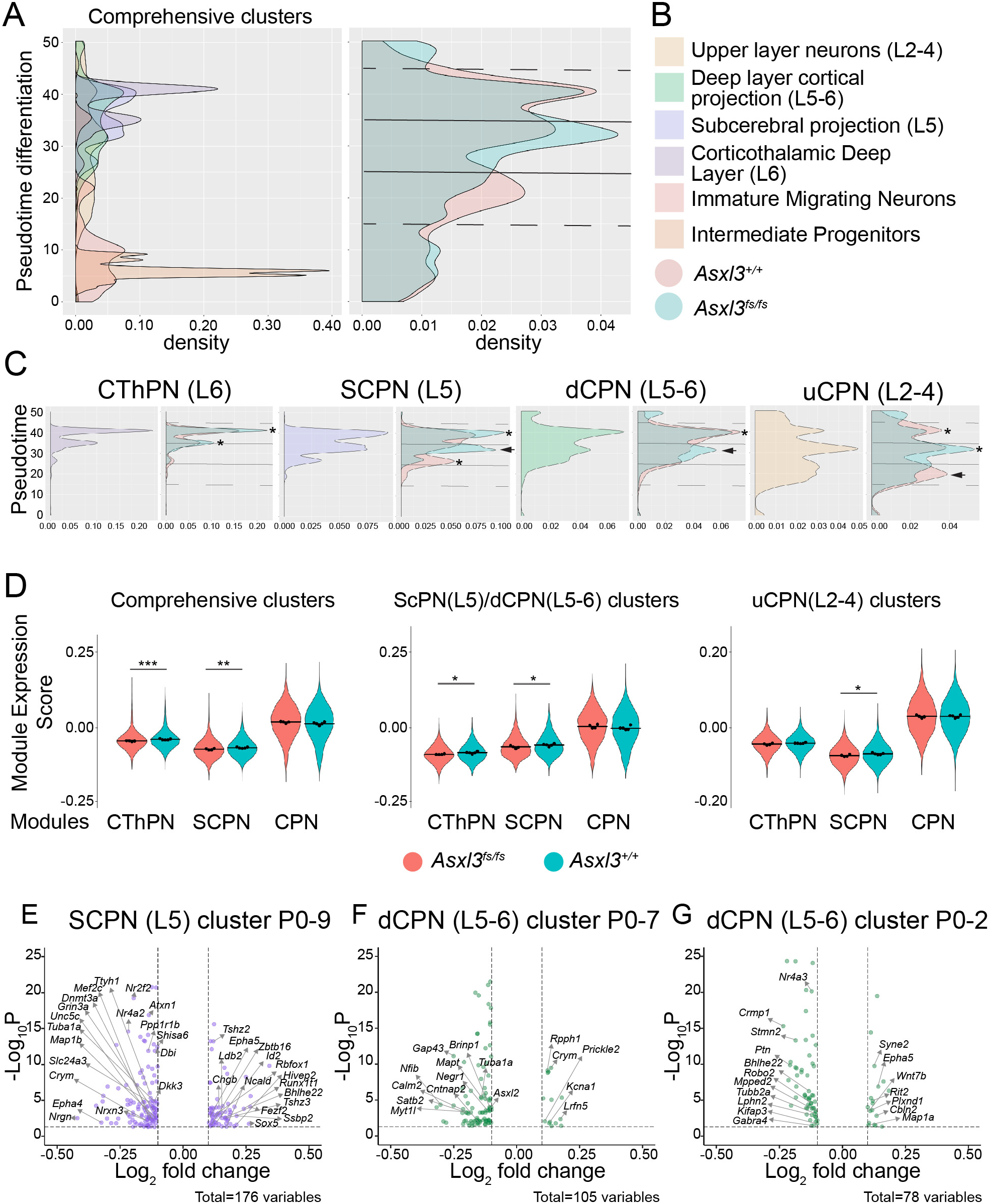
Timing of deep layer neuron differentiation disrupted in *Asxl3^fs/fs^* cortical development A, Histograms of the number of P0.5 cells ordered along pseudotime colored by cell type (left) and colored by genotype (right: *Asxl3^+/+^*, red; *Asxl3^fs/fs^*, blue). **B,** Major cell types and corresponding color code **C,** Individual P0.5 pseudotime histograms of CThPN, SCPN, dCPN, and uCPN. An asterisk indicates peaks with changes in amplitude and arrowheads indicate peaks with a shift in pseudotime. **D,** Violinplots showing module scores for CThPN, SCPN, and CPN gene modules identified by Molyneaux et al. 2015. Module scores were calculated for comprehensive clusters (CThPN, SCPN, dCPN, uCPN), layer 5 neurons (SCPN, dCPN), and upper layer neurons (uCPN). Volcanoplots showing differentially expressed genes identified for **E,** Cluster P0-9 SCPN, **F,** Cluster P0-7 dCPN, and **G,** Cluster P0-2 dCPN.

Pseudotime trajectories for individual cortical neuronal subtypes were generated to identify the contribution of individual neuron classes to genotypic differences in differential peak dynamics of the combined data (Fig. 6C). The independent trajectories of terminally differentiated neuron subtypes do not follow a discrete tip-to-tail order along the pseudotime axis and instead exhibit overlapping distribution. The relative distribution along pseudotime is positively correlated to timing of terminal fate acquisition, with the uCPN L2-4 trajectory encompassing the largest portion of pseudotime (10-50), followed by dCPN (15-50) and SCPN (25-45), with CThPN L6 sharing the smallest portion of the pseudotime axis (30-45) (Fig. 6C). Several genotypic differences in uCPN, SCPN and dCPN classes of neurons contribute to the *Asxl3^fs/fs^* amplitude increase between 25-35. For *Asxl3^fs/fs^* uCPN within this pseudotime interval, we find a differential increase in peak amplitude compared to control. While in the SCPN and dCPN individual trajectories, novel asynchronous *Asxl3^fs/fs^* peaks were detected between the pseudotime 25-35 interval, that do not correspond to smaller *Asxl3^+/+^* peaks of the same periodicity across pseudotime (Fig. 6C). The unique P0.5 *Asxl3^fs/fs^* dCPN and SCPN peaks corresponds to the shift in E14.5 peak periodicity across pseudotime differentiation, further implicating delayed differentiation and altered mature excitatory neuron composition in *Asxl3^fs/fs^* cortex.

To characterize the transcriptional differences underling these altered fate trajectories, the DeCoN corticogenesis transcriptomic resource was used to compile modules of proneural genes which inform CThPN, SCPN and CPN fate acquisition (Table S1) [48]. Differential module expression was assessed for all clusters combined, as well as clusters that broadly comprise deep layer neurons (CThPN; P0-6, SCPN; P0-9 & dCPN; P0-2, P0-7) and upper layer uCPN (P0-0, P0-3, P0-4, P0-5, P0-8) clusters (Fig. 6D). SCPN module expression was significantly increased in *Asxl3^fs/fs^* samples for each cluster class evaluated (adjusted *p*=0.009 comprehensive clusters; *p*=0.030 deep layer clusters; *p=*0.048 uCPN clusters, linear mixed models). Expression of the CThPN module was increased in all clusters (adjusted *p*= 0.002) and deep layer clusters (adjusted *p* = 0.028) (Fig. 6D), while the CPN module was not significantly differentially regulated in any of the cluster groups.

We identified the differentially expressed genes for clusters P0-9, P0-7 and P0-2, to explore the *Asxl3^fs/fs^* transcriptomic changes that drive asynchronous SCPN (cluster P0-9) and dCPN (cluster P0-7 & cluster P0-2) peaks in individual neuronal subtype trajectories. DEGs associated with asynchronous SCPN and dCPN pseudotime differentiation included *Bhlhe22, Stmn2, Fezf2, Ldb2, Tshz2, Crym* and *Cbln2, Plxnd1, Robo2, Ptn, Runx1t1, Ssbp2, Shisa6, Nr2f2, Nr4A2, Myt1l* respectively (Fig. 6E-G). Dysregulation of these genes which drive identity and specification of excitatory lineages, implicates that transcriptomics were impacted by delayed timing of cortical neuron differentiation during *Asxl3^fs/fs^* corticogenesis.

### ASD genes differentially expressed in *Asxl3^fs/fs^* cortex

Large scale human genetic studies have identified hundreds of ASD risk genes, *de novo* mono-allelic frameshift variants in *ASXL3* among them [10-12]. Differential expression of other candidate ASD risk genes has repeatedly been observed in transcriptomic studies of human and mouse ASD neuropathology [32, 49-54]. To identify ASD biology implicated by and shared with biallelic *Asxl3^fs/fs^* molecular pathology, we identified ASD genes differentially expressed in *Asxl3^fs/fs^* transcriptomics. E13.5, E14.5, and P0.5 cluster specific DEGs were intersected with the SFARI ASD gene set, and further partitioned based on high-confidence classification (hcASD) (Fig. 7) [10]. Greater than 12% of all DEGs at each time point were ASD risk genes, which were found to be up- and down-regulated, in a developmental and cell-type defined manner (Fig. 7A-C). At E13.5 *Asxl3^fs/fs^* DEGs are predominantly down-regulated, while at E14.5 and P0.5 differential expression is more equally up- and down-regulated. This is consistent with ASXL3-dependent role in ASD gene expression and provides further evidence for convergence of ASD risk genes on a common molecular mechanism.

**Fig. 7.**
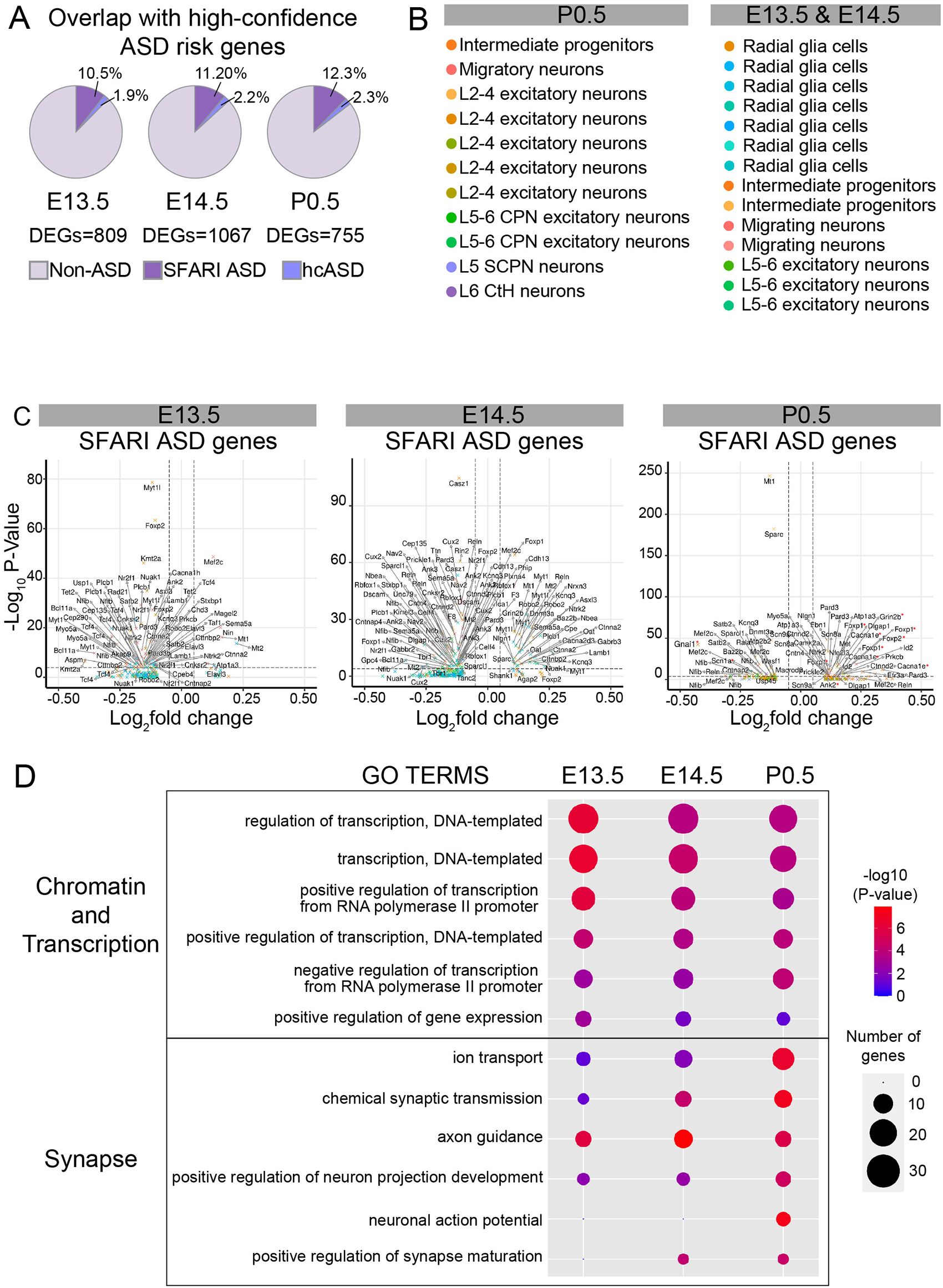
Enrichment of ASD risk genes A, The pie chart illustrates the percentage of SFARI ASD genes and high confidence ASD genes (Satterstrom et al. 2020) amongst our P0.5, E14.5, and E13.5 differentially expressed genes. **B,** Cell type annotations and color codes used for volcano plots in C. **C,** Volcano plots for differentially expressed SFARI ASD genes in E13.5, E14.5, and P0.5 excitatory cells. Genes with a *p*-value < 0.05 and log fold change > 0.1 are displayed and colored by the cell type they are dysregulated in. **D,** Gene ontology analysis showing statistically significant enrichment for dysregulated ASD genes in the E13.5, E14.5, and P0.5 cortex from C. At early timepoints GO terms were enriched for chromatin ASD genes. At later timepoints GO terms were enriched for synaptic ASD genes. The size of each dot represents the number of genes and the shading represents the -log_10_ *p*-value.

Genes that encode proteins involved in chromatin regulation, transcriptional regulation and synaptic signaling have been identified as the genetic basis of ASD [12, 55]. With little evidence of functional overlap between ASD genes, a common molecular pathway is not clear. To investigate functional overlap between DEGs, GO analysis of ASD DEGs at individual developmental time points was performed. Top GO terms identified at E13.5 highlight the function of ASD chromatin and transcriptional regulator genes early in development (Fig. 7D). At E14.5 GO terms reflective of synaptic function and chromatin biology are enriched (Fig. 7D). By P0.5, GO terms associated with synaptic function are significantly enriched. These results implicate molecular synergy during development where chromatin genes are required for early stages of corticogenesis, that ultimately determine synaptic functions at later stages (Fig. 7D). Future experiments will be required to determine if chromatin genes such as *Asxl3* impact synaptic function by directly modulating synaptic genes or indirectly by initiating defects in corticogenesis that disrupt synaptic formation and function. Understanding the convergence of ASD risk genes could provide opportunities to develop targeted therapeutic approaches.

## DISCUSSION

Polycomb transcriptional regulation is critical for organ development and cell fate specification. Evaluation of an *Asxl3* mouse model revealed a critical role in cortical development. The emerging pathogenic model based on analysis of multiple cell types, at sequential developmental timepoints, implicates overactivation of Notch signaling that alters NPCs proliferation and timing of differentiation. These early developmental defects lead to altered composition of excitatory neurons with aberrant expression of proneural genes responsible for layer specificity at later timepoints. Across cortical development, dysregulated genes were enriched for high confidence ASD risk genes, implicating a convergent pathological mechanism. Together our findings underscore the importance of ASXL3 in Polycomb transcriptional repression during development and provide insight into developmental mechanisms altered by ASD risk genes.

### Polycomb transcriptional repression required for corticogenesis

The role of H2AUb1 in Polycomb transcriptional repression remains poorly understood compared to other histone modifications. Pathogenic variants in genes which encode individual components of both PRC1 and PR-DUB are the genetic basis for a growing number of neurodevelopmental disorders, highlighting the importance for dynamic exchange of H2A ubiquitination in neural development [18, 56, 57]. An ASXL3-dependent developmental and transcriptomic disruption of excitatory neuron lineage differentiation and fate was identified in the *Asxl3* frameshift mouse model, with IHC revealing a 25% reduction of BCL11B L5 neurons (Fig. 1). This stands in contrast to a 30% increase in BCL11B cells described in the cortex after conditional knockout of *Ring1b*, the enzymatic component of PRC1 [15]. The opposing lamination defects of these two mouse models mirrors the antagonistic activity of PRC1 and PR-DUB, further implicating a role for dynamic H2AUb1 regulation in NPC fate specification. Future studies need to determine targets of PR-DUB complexes that allow for dynamic transcriptional control of proneural genes.

### Extrinsic signaling pathways coordinate balance of self-renewal & differentiation

During corticogenesis NPCs undergo symmetric self-renewing and asymmetric neurogenic mitotic events. Extrinsic signaling pathways, including Notch, Wnt, and Hippo, are important for directing the self-renewal and differentiation behavior of NPCs. Overactivation of Notch or Wnt signaling or inactivation of Hippo signaling during neurogenesis shift NPCs toward self-renewal and away from neurogenic proliferation [37, 39, 41, 42]. A commonly observed phenotype in these mouse models is lateral expansion of RGCs and increased VZ perimeter, without a corresponding change in brain size or cortical thickness, similar to the phenotypes observed in *Asxl3^fs/fs^* brain development [37, 39, 41, 42]. Notably, the extent of lateral expansion was shown to be dose-dependent implicating a critical role for ASXL3 dependent transcriptional regulation of extrinsic signaling pathways as *Asxl3^fs/fs^* transcriptomic changes implicated their dysregulation at an early stage of neurogenesis [37, 41, 58].

The N1ICD directly regulates expression of WNT and Hippo signaling pathways components, allowing Notch to facilitate pathway crosstalk in NPCs [37]. Consistent with altered Notch signaling, *Hes1* and *Hes5* are up-regulated in *Asxl3^fs/fs^* mice. These transcription factors are at the nexus of the Notch signaling pathway during corticogenesis, required for inhibiting neuronal differentiation and promoting maintenance of NPC multipotency [59-62] Phenocopy between *Asxl3^fs/fs^* and the transgenic *Hes1*- and *Hes5*-overexpression mouse models, implicates a prominent role for Notch signaling through HES1/HES5 in *Asxl3^fs/fs^* neuropathology [38, 43]. In both models, *Hes1* or *Hes5* overexpression results in lateral expansion of VZ RGC progenitors and altered emergence of excitatory fated cortical neurons. HES1 and HES5 maintain NPCs multipotency by repressing *Ngn2*, which induces neurogenesis. Using the cell-type and temporal resolutions of Asxl3^fs/fs^ transcriptomic data, the overexpression of HES1 and HES5 in *Asxl3^fs/fs^* RGCs at E13.5 was correlated to differential expression of proneural genes, including down-regulation of *Ngn2* in *Asxl3^fs/fs^* differentiating cell types at E14.5 [38, 43]. Ultimately, the developmental impact on corticogenesis can be summarized into a model where NPC self-renewal alters the length of neurogenesis through lateral inhibition which affects the fate of excitatory neuron and cytoarchitectural cortical layer deposition [38, 43].

### Single-cell transcriptomic view of cortiocogenesis

Histology based analysis of corticogenesis has identified individual proneural genes critical for generation and differentiation of distinct cortical neuron subtypes [63-67]. Generating the complement of mature cortical neuron subtypes from multipotent NPCs requires cell-intrinsic and dynamic chromatin modifications that play important roles in defining transcriptional plasticity and cell identities. Distinguishing the transcriptomic diversity of individual cells in a developing organ requires single-cell resolution. The challenge moving forward is to correlate single-cell transcriptomic characterization of normal and pathogenic corticogenesis with decades of knowledge generated by investigating corticogenesis in single-gene mouse models using a limited selection of histological markers. This correlation is less complicated at earlier embryonic timepoints when cortical tissue is largely composed of RGCs and IPs expressing consistent marker genes that modulate minimally during corticogenesis. At later stages of corticogenesis, transcriptional programs of excitatory lineages diverge in the post-mitotic neurons as cortical neurons separate into increasingly partitioned cell populations.

Due to the relatively low heterogeneity of E13.5 and E14.5 cortical tissue, we observed high correlation between scRNA-seq data and tissue analysis with immunohistochemistry. While the ratio of progenitors detected with scRNA-seq decreased between E13.5 and E14.5, the progenitor marker genes were consistent between stages and were robustly distinguished from terminal neuronal subtypes. The conformity between these two techniques allowed us to predict the Asxl3^fs/fs^ developmental mechanisms based on the transcriptomics. Subsequent evaluation of the cell-specific transcriptomics *in vivo* identified important molecular mechanisms of early *Asxl3^fs/fs^*neuropathology.

Greater diversity of terminally differentiated neuronal subtypes at P0.5 complicates correlating the scRNA-seq data with immunohistochemistry. Since we detect a reduction in BCL11B L5 neurons with immunohistochemistry, we predicted that our scRNA transcriptomic analysis would directly translate into a reduction of SCPN neurons. Instead, we observe a significant shift in the proportions of all neuronal subtypes in *Asxl3^fs/fs^*, consistent with a transcriptional shift in cell lineage profiles that transform their terminal fate. This interpretation is supported by the divergent P0.5 *Asxl3^+/+^* and *Asxl3^fs/fs^* pseudotime differentiation peaks. The upregulation of CThPN and SCPN cell lineage profiles resembles transcriptional alterations attributed to Polycomb transcriptional regulation that result in fate transformations [16, 36]. In addition, the dCPN and SCPN DEGs we detect for individual trajectories include proneural genes known to govern the acquisition of cell identity. These data are consistent with cortical neuron fate changes in *Asxl3^fs/fs^* samples that acquire a hybrid transcriptional profile, which may allow these cells to associate with clusters across the SPCN, dCPN and uCPN classes. The broad distribution of dCPN and uCPN classes of neurons mapped to a spatial reconstruction of the visual cortex may reflect the current nuances in the annotation of cortical classes of cells by transcriptional profile, as opposed to detection of protein expression with immunohistochemistry. In summary, a reduction in BCL11B L5 neurons is detected by IHC in *Asxl3^fs/fs^* cortex, without a change in L5 cell density. This suggests these cells are replaced by another neuron type or an undefined hybrid fate that may reflect overlapping expression of CThPN and SCPN proneural genes based on transcriptomic data.

### Molecular synergy of ASD neuropathology

Our data linking early defects in *Asxl3^fs/fs^* cortical neurogenesis to altered timing of cortical neuron subtypes is pertinent in the context of human ASD genetics and *in vitro* models that predicted pathogenic mechanisms that impinge on neurogenesis [54, 68-74]. Recently, neural fated tissue differentiated from human pluripotent stem cells (hPSCs) carrying pathogenic variants in hcASD genes were tested for neurogenesis competency. Consistent with the *Asxl3^fs/fs^* neuropathology, the cellular progeny of *ASXL3* hPSC lines fell into a subclass of ASD risk genes that are hyporesponsive to WNT and exhibit decreased neuronal output with an enrichment for NPCs [68]. In the same phenotypic class of ASD risk genes were the chromatin associated genes *ASH1L*, *KDM5B*, *CHD2* and *KMT2C* that have a documented role in impacting Polycomb transcriptional regulation and implicate novel molecular mechanisms of ASD yet to be explored in animal models. Conversely, neurogenesis defects which result in increased neurogenesis at the expense of SOX2 progenitors were observed in *Bcl6* gain-of-function mouse model that were in part due to dysregulation of extrinsic signaling pathways. BCL6, a transcriptional repressor that interacts with the PRC1.1 component, BCOR, that modulates H2AUb1 [75]. Together these findings demonstrate that dynamic Polycomb transcriptional repression is essential for balancing progenitor expansion and differentiation of NPC.

ASD neuropathology is attributed to mid-gestation neurogenesis defects in the prefrontal cortex and excitatory/inhibitory (E/I) imbalance of the cortical microcircuit [76-78]. *Asxl3^fs/fs^* neuropathology provides evidence for how pathogenic variants in chromatin genes which disrupt development could damage synaptic function and the balance of cortical microcircuit activity. Such imbalance could be initiated due to a change in the composition of mature cortical subtypes. The shift in the focus of GO terms from chromatin and transcriptional regulation at E13.5 to greater enrichment in synaptic function terms at P0.5 support this interpretation. Precluding direct differential expression of synaptic genes by chromatin based genetic etiologies of ASD, *in vivo* studies have demonstrated that even subtle developmental alterations in circuit composition can result in disproportionally large functional effects on the mature circuit [79-81].

## METHODS

### Animals

All experiments were performed in accordance with animal protocols approved by the Unit for Laboratory Animal Medicine (ULAM), University of Michigan. The Asxl3 null mice line was generated by cloning sgRNAs that target a region in exon 12 into a pX330 vector. The vectors were microinjected into fertilized eggs before being transferred into pseudopregnant C57BL/6 X DBA/2 F1 females. Topo cloning and sanger sequencing were used to detect a unique frameshift *Asxl3* allele. *Asxl3^+/fs^* mice were maintained on a C57BL/6 background. Heterozygous breeding was used for experiments with E0.5 established as the day of vaginal plug.

### Western Blot Analysis

Whole brains from E13 *Asxl3^+/+^* and *Asxl3^fs/fs^* mice were homogenized in RIPA buffer supplemented with protease inhibitor cocktail and phosphatase inhibitor cocktail 3 obtained from Sigma-Aldrich (P8340 and P0044; St Louis, MO, USA). Protein levels were normalized after BCA analysis (Pierce). Cell lysates were separated using electrophoresis on 4-20% SDS-polyacrylamide gels and transferred to PVDF membrane (Millipore, Billerica, MA, USA). For western blot, after the transfer, the PVDF membrane was blocked with 5% milk and incubated with following antibodies overnight. Primary antibodies used were: anti-ASXL3 (Bielas Lab, 1 to 200), anti-ubiquityl-Histone H2A (Cell Signaling Technology, 8240, 1 to 2000), anti-Histone H3 (Abcam, Ab10543, 1 to 5000). Donkey anti-rabbit HRP-conjugated (Cytiva, NA9340V, 1 to 5000) and goat anti-mouse HRP-conjugated (Invitrogen, 32430, 1 to 10000) were used for 1h incubation at room temperature. Antibody incubation and chemiluminescence detection were performed according to manufacturer’s instruction [ThermoFisher Scientific, Waltham, MA, USA, cat no. 34095].

### Immunohistochemistry and cell counting

Brains were dissected and removed from mice at E13.5, E14.5, E15 or P0 and then kept in 4% PFA at 4°C overnight. Brains were cryopreserved by submersion in 20% then 30% sucrose solutions. After embedding in OCT cryosectioning media (Tissue-Tek, Torrance, CA), brains were cryosectioned at 13 μm. Sections were incubated with PBS for 15 min to wash away OCT. For antibodies that required antigen retrieval, cryosections were heated in 10 mM Sodium citrate for 20 minutes at 95°C followed by incubation at room temperature for 20 minutes. Incubation with a normal donkey serum blocking buffer [5% NDS (Jackson ImmunoResearch), 0.1% Triton X-100, 5% BSA]. was performed for 1 hour. Sections were stained with primary antibodies in blocking buffer at 4℃ overnight, washed with PBS, and stained with secondary antibodies at room temperature for 1 hour. Slides were washed with PBS, incubated with DAPI for 5 minutes and cover-slipped with MOWIOL. Images were acquired with a Nikon A1 microscope and processed with LAS X software. Cells were counted using ImageJ. The following antibodies and dilutions were used: SOX2 (Neuromics, GT15098, 1:500), CTIP2 (Abcam, ab18465, 1:500), TBR1 (Abcam, ab31940, 1:200), SATB2 (Abcam, ab51502, 1:100), CUX1 (Santa Cruz, sc-13024, 1:150), pH3 (Abcam, ab10543, 1:250), EOMES (Abcam, ab23345, 1:400). AlexaFluor-conjugated secondaries were: donkey anti-rabbit 647 (Invitrogen, A31573, 1:400), anti-rabbit 555 (Invitrogen, A31572, 1:400), donkey anti-mouse 555 (Invitrogen, A31570, 1:400), donkey anti-mouse 647 (Invitrogen, A31571, 1:400), donkey anti-goat 555 (Abcam, ab150154, 1:400), donkey anti-goat 488 (Invitrogen, A-11055, 1:400).

### EdU Birthdate analysis

Time pregnant dames were injected with EdU (20 mg/kg) at embryonic day 13.5 or 14.5 (E13.5 or E14.5). At P0.5, brains were dissected and removed from EdU injected pups and kept in 4% PFA at 4℃ overnight. EdU was labeled and detected in Cryosections with the Click-IT EdU imaging kit (Invitrogen, Carlsbad, CA) according to the manufacturer’s instructions. After sections were incubated with Click-IT reaction cocktail, they were washed with normal donkey serum blocking buffer. Then, additional antibody staining was performed.

### Cresyl Violet

Brains were dissected and removed from mice at P0.5 then kept in 4% PFA overnight. Brains were dehydrated in a series of ethanol washes and embedded in paraffin with a paraffin tissue processor. 6 μm microtome sections were dewaxed in xylene, rehydrated in ethanol, stained with 0.5% cresyl violet, dehydrated with ethanol and cleared with xylene.

### Collection of cells for single-cell RNA sequencing

The cerebral cortices from E13.5, E14.5 or P0.5 brains were dissected in Earl’s balanced salt solution (EBSS). Isolated cortices were incubated with papain (Worthington Biochemical Corporation),5.5 mM L-cysteine-HCL, 1.1 mM EDTA, and 100 mg/ml DNase I in O_2_:CO_2_ equilibrated EBSS for 8 minutes at 37℃. Samples were titrated with flame tipped glass pipettes and centrifuged at 800 RCF for 5 minutes. Cells were resuspended with ovomucoid protease inhibitor (Worthington Biochemical Corporation) and 50 mg/ml DNase I in O_2_:CO_2_ equilibrated EBSS and passed through a 70 micron cell strainer. Cells were centrifuged at 200 RCF for 5 minutes over a gradient of ovomucoid protease inhibitor and cells were resuspended in N2/B27 media. Cells were counted and checked for viability using a Luna automated cell counter before loading onto a Seq-Well platform.

### Seq-Well Single Cell RNA-sequencing

Seq-Well was performed as described [31, 82]. Briefly, functionalized Seq-Well arrays, containing 90,000 picowells, were loaded with barcoded beads (ChemeGenes, Wilmington, MA). 20,000 cells were loaded onto the arrays and incubated for 15 minutes. To remove residual BSA and excess cells, arrays were washed with PBS. Functionalized membranes were applied to the top of arrays, sealed in an Agilent clamp, and incubated at 37 for 45 minutes. Sealed arrays were incubated in a lysis buffer (5M guanidine thiocynate, 1mM EDTA, 0.5% sarkosyl, 1% BME) for 20 minutes followed by a 45 minute incubation with hybridization buffer (2M NaCl, 1X PBS, 8% PEG8000). Beads were removed from arrays by centrifuging at 2000xg for 5 minutes in wash buffer (2M NaCl, 3mM MgCl_2_, 20mM Tris-HCl pH 8.0, 8% PEG8000). To perform reverse transcription beads were incubated with the Maxima Reverse Transcriptase (Thermo Scientific) for 30 minutes at room temperature followed by overnight incubation at 52℃. Reactions were treated with Exonuclease 1 (New England Biolabs, M0293S) for 45 minutes at 37℃. Whole transcriptome amplification was performed using the 2X KAPA Hifi Hotstart Readymix (KAPA Biosystems, KK-2602). Beads were split to 1,500-2,000 per reaction and run under the following conditions 4 Cycles (98℃, 20s; 65℃, 45s; 72℃, 3m) 12 Cycles (98℃, 20s; 67℃, 20s; 72℃, 3m) final extension (72℃, 3m, 4℃, hold). Products were purified with Ampure SPRI beads (Beckman Coulter, A63881) at a 0.6X volumetric ratio then a 1.0X volumetric ratio. Libraries were prepared using the Nextera XT kit (illumina) and libraries were sequenced on an Illumina NextSeq 75 cycle instrument.

### scRNA-seq data processing

Sequencing reads were processed into a digital gene expression matrix using Drop-seq software as described [83]. FASTQ files were converted into bam files before being tagged with cell and molecular barcodes and trimmed. After converting back to FASTQs, reads were aligned to mm10 with STAR. BAM files are then sorted, merged, and tagged with gene exons. Bead synthesis errors were corrected as described and digital gene expression matrices were generated. We excluded the poor-quality cells in the gene-cell data matrix using the Seurat package (v3.1.2) for the samples (E13.5: n=8, E14.5: n=9 and P0.5: n=9). For downstream analysis cells with fewer than 300 detectable genes, greater than 5,000 genes or greater than 10% mitochondrial genes were removed. Genes that were detected in less than 5 cells were removed. To remove blood cells from our data we visualized the raw count of hemoglobin genes (*Hbb-bh1*, *Hba-a2*, *Alas2*, *Hbb-y*, *Hbb-bs*, *Fam46c*, *Hbb-bt*, *Hba-a1*, *Hmbs*, *Fech*, *Slc25a37*) using violin plots to set a threshold for filtering blood cells. We excluded the cells that expressed more than these predetermined thresholds for further analysis. We normalized the data in each sample for each cell by dividing the total counts for that cell and multiplied by 10,000. This is then natural-log transformed using log1p. After this normalization step, we selected genes showing a dispersion (variance/mean expression) larger than two standard deviations away from the expected dispersion as variable genes using the Seurat (v3.1.2) function “FindVariableGenes” for each sample. In this way, there are 2000 significant genes which were identified using variance stabilizing transformation for each sample in different stages. We then used these most informative genes for sample alignment for each stage. We integrated all the samples in each stage by using the FindIntegrationAnchors and IntegrateData functions of Seurat (v3.1.2). We performed this alignment as described and implemented in Seurat tutorial (v3.1.2). After we integrated the datasets for each stage, we scaled and centered the data for each stage.

### Dimensionality Reduction, Visualization and Clustering

We used the Seurat package (v3.1.2) to perform dimensionality reduction. We used the integrated and normalized data as the input to the RunPCA function of Seurat (v3.1.2) in order to compute the first 100 PCs. After that, we used the elbow algorithm to find the optimal number of PCs to construct Uniform Manifold Approximation and Projection (UMAP) plots. Visualizations in a two-dimensional space were done using RunUMAP and RunTSNE function of Seurat (v3.1.2) for the integrated data using previous dimensional reduction data and predetermined best PC number. We performed a graph-based clustering approach using FindNeighbors and FindClusters functions of Seurat (v3.1.2). We clustered the cells based on modularity optimization technique: Louvain algorithm with resolution parameter 0.5 and partitioned the graph constructed before into communities. We then collected cluster marker genes using the Wilcoxon rank-sum test between the cells in a single cluster and all other cells with log fold change threshold of 0.2. To assign identities to clusters, we cross-referenced the marker genes with previously described cortical cells [29, 30, 47]. The expression of cell type markers was visualized using the FeaturePlot function in Seurat. After we determined the identity of excitatory and non-excitatory cortical cells, we extracted the cell types that contribute to the excitatory cells and redid the clustering for this subset.

### Alignment with Loo et al. 2019

We mapped our dataset to the Loo et al. 2019 [29] E14.5 and P0 cortices using label transferring features from Seurat (v3.1.2). We leveraged the cell type labels provided by Loo et al. 2019 as the reference and projected our cells onto the reference using FindTransferAnchors and TransferData function from Seurat. We obtained the classification scores which depicts the level of cell type prediction for the query cells and assigned the reference labels to the query cells based on the highest prediction score. For our excitatory neuron lineage analysis, we filtered out cells that do not contribute to this lineage after we projected the reference labels on our dataset. P0 filtered cells included: Astrocytes (10-P, 13-P), Choroid Plexus (20-P), Endothelial (17-P, 21-P), Ganglionic Eminence (9-P), Interneurons (5-P,14-P, 11-P, 6-P), Layer 1 (19-P), Microglia (22-P), Oligodendrocytes (16-P), Striatal neurons (3-P, 7-P). E14.5 filtered cells included: Choroid Plexus (22-E), Endothelial (18-E), Ganglionic Eminence (6-E), Interneurons (1-E, 12-E), Microglia (20-E), Cortical Hem RGC (21-E), Striatal neurons (16-E, 9-E), Thalamic (19-E). We visualized the query transferred labels on the UMAP plots using DimPlot function from Seurat.

### Differential expression analysis and enrichment analysis

We used the Wilcoxon ran-sum test to conduct the DE analysis between *Asxl3^+/+^* and *Asxl3^fs/fs^* samples for each stage in the alignment data. To investigate the overlap of ASD risk genes with DEGs within individual clusters, we overlapped the combined list of DEGs at each timepoint with genes listed in the SFARI database (https://gene.sfari.org/, 02-11-2020 release). We combined all the significant genes from the DE analysis between *Asxl3^+/+^* and *Asxl3^fs/fs^* samples for each stage in the alignment data across different clusters. We used the ‘EnhancedVolcano’ R package to visualize ASD associated DEGs with a log fold change greater than 0.05 and p-value less than 10e-5. To investigate the overlap of Notch target genes, we evaluated the enrichment of direct and indirect N1ICD targets identified by Li et al. 2012 [37] by conducting a Fisher’s exact test on the 2×2 enrichment table (DAVID). We utilized the cluster specific DE gene list and compared it to the direct and indirect N1ICD gene lists to collect the 2×2 enrichment table which was then evaluated by the two-sided Fisher exact test. We used the p-value collected from Fisher exact test and the number of genes that are contained in the gene list in each cluster to visualize the enrichment results on each cluster across different stages.

### Functional Enrichment Analysis

We used lists of significant genes from differential expression analysis across genotypes to identify enriched biological pathways using the Database for Annotation, Visualization and Integrated Discovery (DAVID) v6.8, an online program uses a modified Fisher’s exact test (EASE score) to examine the significance of gene-term enrichment. Gene names were first mapped into Entrez ID through genome wide annotation for mouse by R package ‘org.Mm.eg.db’. KEGG pathway, UniProt keywords and Gene Ontology (GO) were chosen for three different options of annotation categories. Terms were filtered using a threshold of Bonferroni corrected P-value smaller than 0.05. GO terms including molecular function, cellular component and biological process were also used for DAVID functional annotation clustering, with default classification stringency.

### Module Score

To investigate the differential expression pattern of cortical neuron subtype modules, we used the gene list from Molyneaux et al. 2015 and utilized the ‘AddModuleScore’ function using the Seurat package (v3.1.2). For CThPN we used genes in cluster 6 and 9, for SCPN we used genes in cluster 16, and for CPN we used genes in cluster 0, 5, and 15. For each module, the average gene expression level was calculated and was compared to the control genes for each cell. Moreover, we used a linear mixed-effect model to evaluate the module expression differences between *Asxl3^+/+^* and *Asxl3^fs/fs^*. Specifically, we treated the module scores as the output for each cell, genotype as the predictor variable with the sample identity being the random intercept to account for the batch effect. We applied the ANOVA test to calculate the p-values which are then adjusted across modules using the Benjamini-Hochberg correction.

### Spatial Distribution

To learn the spatial structure of P0 data, we used Tangram [45] and aligned our data with STARMap dataset [46] (1127 cells and 166 genes). We manually assigned the spatial clusters to the cells based on the neuronal layer specific genes and used the ‘shiny’ package to label the cells. We then extracted the common genes among our data and the STARMap data and obtained the cell-by-coordinates structure matrix where it gives the probability for cell to be in coordinates. The matrix was then multiplied by the one hot encoding matrix of cell types in our data, and we visualized the mapping results using the ‘ggplot2’ package.

### Pseudotime analysis

For neuronal cells identified using two levels of clustering in each stage, we used Monocle 3 to conduct single-cell pseudotime analysis [84]. First, we performed principal components analysis on gene expression data to capture the variation and removed batch effects using mutual nearest neighbor alignment. UMAP was then used for dimensionality reduction and visualization, setting the number of neighbors to 50. We then created a curvilinear trajectory using the function ‘learn graph’ and ordered the cells with progenitors (in E13.5 and E14.5) or intermediate progenitors (in P0) constrained as starting points. The 2-D visualization of trajectory was conducted using default function ‘plot cell’ in Monocle 3 and the comparison of pseudotime information between genotypes was shown using density plot and histogram by ggplot2.

## Supporting information

McGrath et al. Supp Data File

## ACKNOWLEDGEMENTS

Work was supported by the Eunice Kennedy Shriver National Institute of Child Health and Human development (R01AWD010411 to SLB), National Institute of Neurological Disorders and Stroke (R01NS101597 to SLB) Simons Foundation Directors Award (19-PAF06833,17-PAF05917 SLB), Leo’s Lighthouse Foundation to SLB and BTM, NIH Cellular and Molecular Biology Training Grant (T32-GM007315 to BTM), Genetics Training Grant (to AM). Department of Biotechnology, Innovative Young Biotechnologist Award (BT/12IIYBAl2019/13) and Ramalingaswami Fellow (<u>D.O.NO.BT/HRD/35/02/2006</u>) to AS.

