## Supplementary material for "ASXL3 controls cortical neuron fate specification through extrinsic self-renewal pathways": McGrath et al. Supp Data File

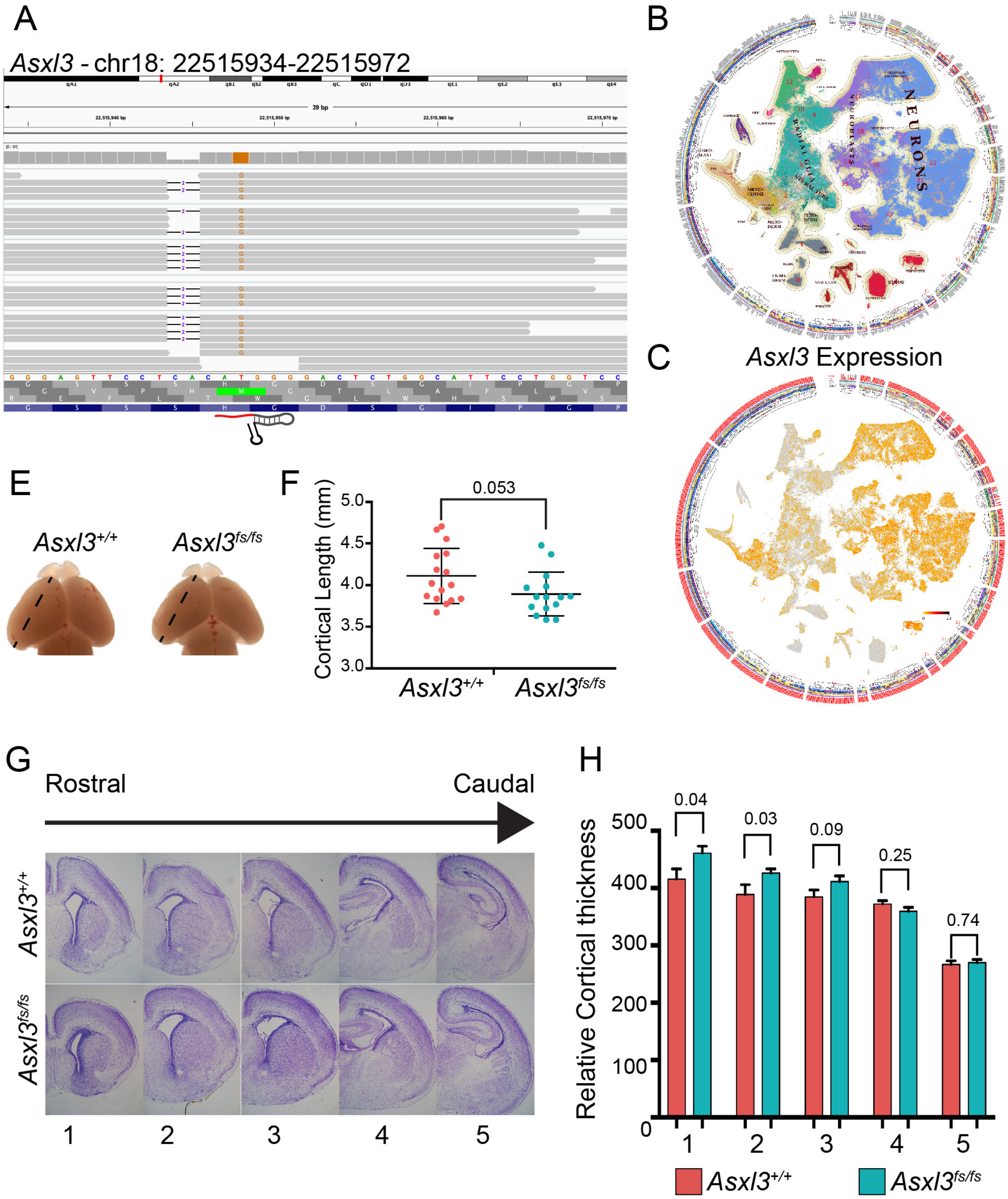
A
Asxl3 - chr18: 22515934-22515972
39 bp
B
NEURONS
NEUROBLASTS
RADIAL GLIA
C
Asxl3 Expression
E
Asxl3+/+
Asxl3fs/fs
F
0.053
Cortical Length (mm)
5.0
4.5
4.0
3.5
3.0
Asxl3+/+
Asxl3fs/fs
G
Rostral
Caudal
Asxl3+/+
Asxl3fs/fs
1
2
3
4
5
H
Relative Cortical thickness
500
400
300
200
100
0
0.04
0.03
0.09
0.25
0.74
1
2
3
4
5
Asxl3+/+
Asxl3fs/fs

**Supplemental Figure 1. Cortical thickness and length unaffected in *Asxl3*$^{fs/fs}$**

**A,** IGV viewer window of *Asxl3* that shows the detection of *Asxl3*$^{fs}$ mRNA in our P0.5 scRNA-seq. **B,** Wheel plots showing scRNA-seq data using tSNE embedding (La Manno et al. 2020) (http://mousebrain.org). Clusters are colored by major classes. **C,** Expression of *Asxl3* overlaid on the tSNE plot in B. **E,** Representative images and **F,** quantification of P0.5 cortical length, marked by a black dashed line, for *Asxl3*$^{+/+}$ ($n$=16) and *Asxl3*$^{fs/fs}$ ($n$=15) mice. $p$=0.053 using two-tailed unpaired Student's $t$ test. **G,** Serial coronal sections of P0.5 brains stained with cresyl violet. Comparable *Asxl3*$^{+/+}$ (top) and *Asxl3*$^{fs/fs}$ (bottom) sections from rostral to caudal regions were compared. **H,** Quantification of cortical thickness in equivalent areas of control and homozygous frameshift cortices (1: $p$=0.24, 2: $p$=0.29, 3: $p$=0.09, 4: $p$=0.17, 5: $p$=0.59 using two-tailed unpaired Student's $t$ test). All values are displayed as mean ± SEM.

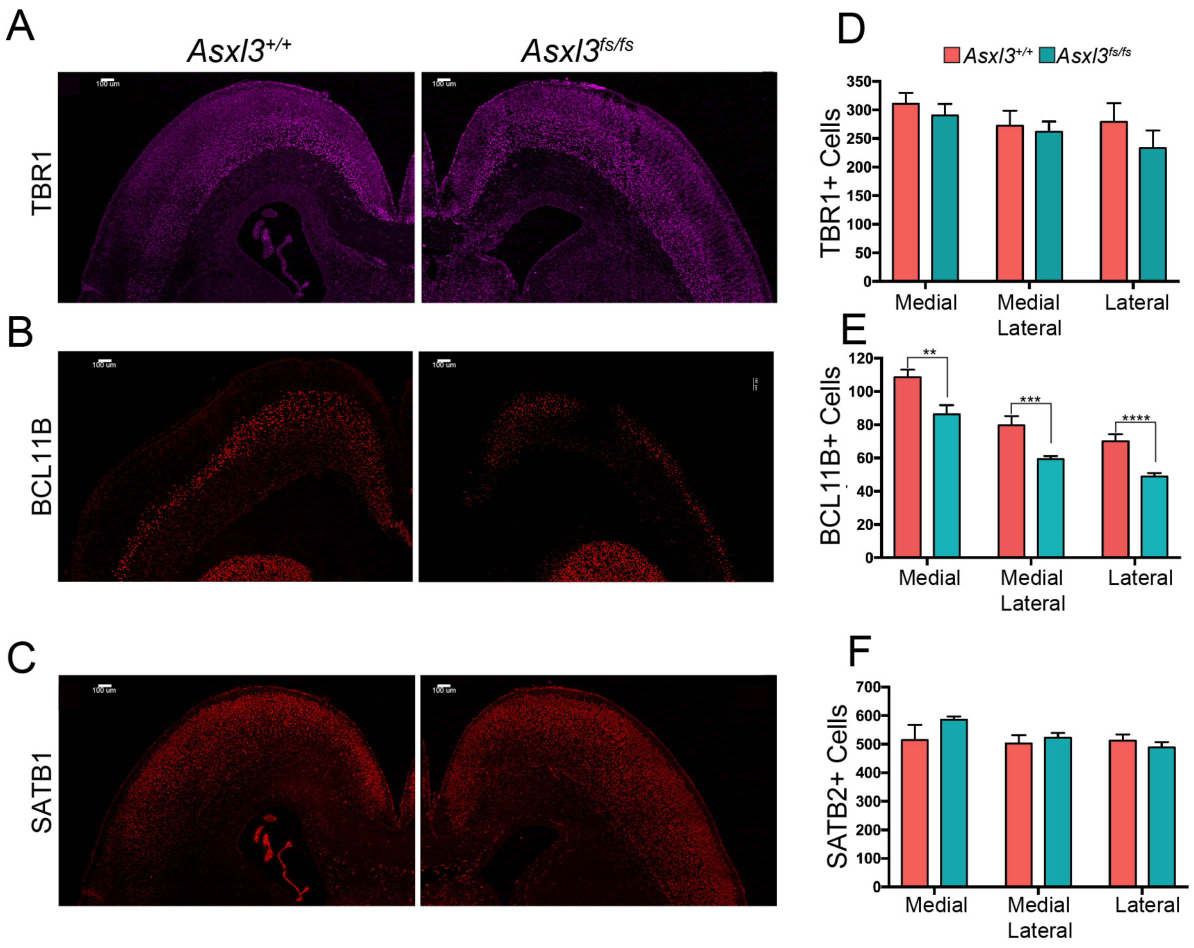

A
$Asxl3^{+/+}$
$Asxl3^{fs/fs}$
TBR1
B
BCL11B
C
SATB1
D
$Asxl3^{+/+}$ $Asxl3^{fs/fs}$
TBR1+ Cells
350
300
250
200
150
100
50
0
Medial
Medial Lateral
Lateral
E
BCL11B+ Cells
120
100
80
60
40
20
0
**
***
****
Medial
Medial Lateral
Lateral
F
SATB2+ Cells
700
600
500
400
300
200
100
0
Medial
Medial Lateral
Lateral

**Supplemental Figure 2. Excitatory neuron cortical composition in *Asxl3*$^{fs/fs}$ mouse model** Large scale images displaying immunostaining of *Asxl3*$^{+/+}$ versus *Asx3*$^{fs/fs}$ P0.5 coronal cortical sections with layer-specific markers **A,** TBR1 (layer 6), **B,** BCL11B (layer 5), and **C,** SATB2 (layer 2-4). Quantification of the number of neurons expressing **D,** TBR1, **E,** BCL11B, or **F,** SATB2 in *Asxl3*$^{+/+}$ ($n$=4, $n$=8, $n$=3) and *Asx3*$^{fs/fs}$ ($n$=4, $n$=8, $n$=3) cortices. Equal-sized bins were used to quantify cells across medial, medial/lateral, and lateral regions. TBR1 (m: $p$=0.49, m/l: $p$=0.76, l: $p$=0.35) BCL11B (m: $p=7.7\times10^{-3}$, m/l: $p=4.3\times10^{-3}$, l: $p=5.3\times10^{-4}$), SATB2 (m: $p$=0.26, m/l: $p$=0.57, l: $p$=0.45) using two-tailed unpaired Student's $t$ test. Scale bars, 100 um. **$p$<0.01, ***$p$<0.005, ****$p$<0.001. All values are displayed as mean ± SEM.

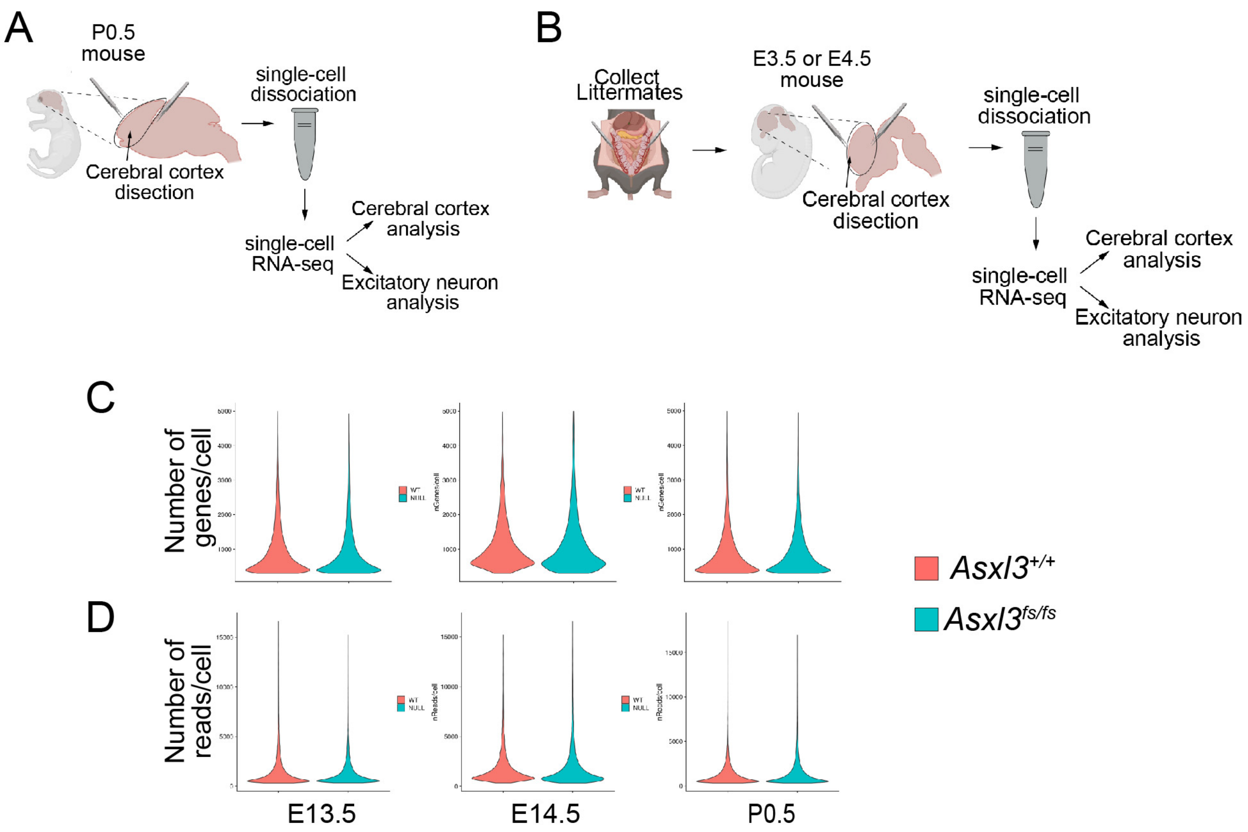


F

| Sample | Genotype | Raw Number of Cells | Final Number of Cells | Number of Reads |
|---|---|---|---|---|
| E13_1 | Asxl3+/+ | 6947 | 3960 | 8271489 |
| E13_2 | Asxl3+/+ | 4220 | 4046 | 3375635 |
| E13_3 | Asxl3+/+ | 6385 | 5370 | 7049063 |
| E13_4 | Asxl3+/+ | 10625 | 6693 | 11573418 |
| E13_5 | Asxl3+/+ | 7679 | 6711 | 10816834 |
| E13_6 | Asxl3fs/fs | 9331 | 8211 | 13355546 |
| E13_7 | Asxl3fs/fs | 6079 | 5237 | 6118518 |
| E13_8 | Asxl3fs/fs | 9734 | 6692 | 10237729 |
| E14_1 | Asxl3+/+ | 2823 | 2290 | 4446278 |
| E14_2 | Asxl3+/+ | 2892 | 2727 | 5773798 |
| E14_3 | Asxl3+/+ | 2546 | 1652 | 1910141 |
| E14_4 | Asxl3fs/fs | 2807 | 2590 | 5994814 |
| E14_5 | Asxl3fs/fs | 2140 | 2016 | 4142246 |
| E14_6 | Asxl3fs/fs | 1540 | 1464 | 3166855 |
| E14_7 | Asxl3fs/fs | 2505 | 1873 | 3390708 |
| E14_8 | Asxl3fs/fs | 4179 | 3462 | 6270291 |
| E14_9 | Asxl3fs/fs | 8904 | 6889 | 11362018 |
| P0_1 | Asxl3+/+ | 10978 | 8459 | 11615000 |
| P0_2 | Asxl3+/+ | 6499 | 3785 | 6599141 |
| P0_3 | Asxl3+/+ | 11067 | 8428 | 11175369 |
| P0_4 | Asxl3+/+ | 3254 | 2839 | 3553872 |
| P0_5 | Asxl3fs/fs | 5853 | 5321 | 8811273 |
| P0_6 | Asxl3fs/fs | 10669 | 9548 | 19413221 |
| P0_7 | Asxl3fs/fs | 13095 | 10077 | 15659975 |
| P0_8 | Asxl3fs/fs | 7608 | 7381 | 7086778 |
| P0_9 | Asxl3fs/fs | 7159 | 6345 | 6776285 |

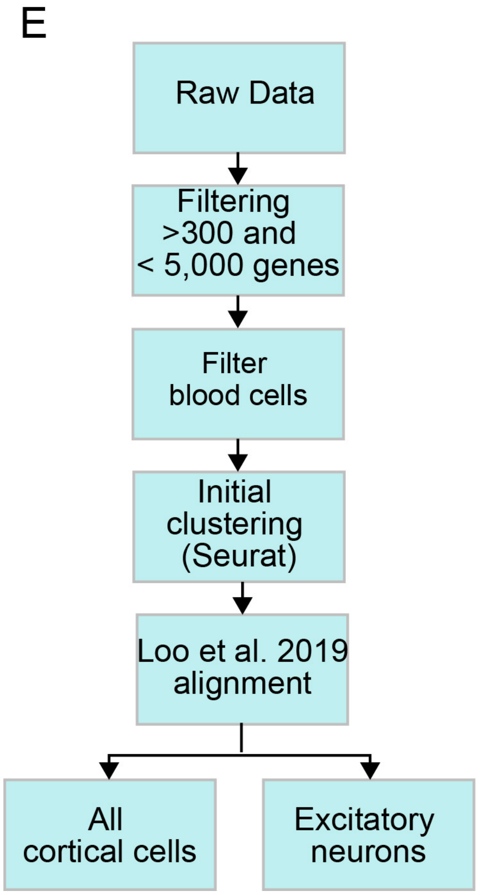

**Supplemental Figure 3. Workflow and QC for scRNA-seq of cortical tissue** Illustration of the **A,** P0.5 and **B,** E13.5/14.5 cortical dissections. Single cell dissociations were loaded onto the SEQ-Well platform and prepped for RNA sequencing. After cell annotation, excitatory neurons were bioinformatically enriched for analysis. Violin plots showing the distribution of per cell **C,** gene and **D,** reads detection at E13.5, E14.5 and P0.5 for *Asxl3*$^{+/+}$ and *Asxl3*$^{fs/fs}$ samples. **E,** Schematic overview of our scRNA-seq analysis pipeline. Cells with fewer than 300 or more than 5,000 genes detected were removed. As were cells expressing blood cell genes. After unsupervised clustering with Seurat, excitatory neurons were selected for further analysis. **F,** Table listing the sample, cell number, number of filtered cells and number of reads for E13.5, E14.5 and P0.5 sets.

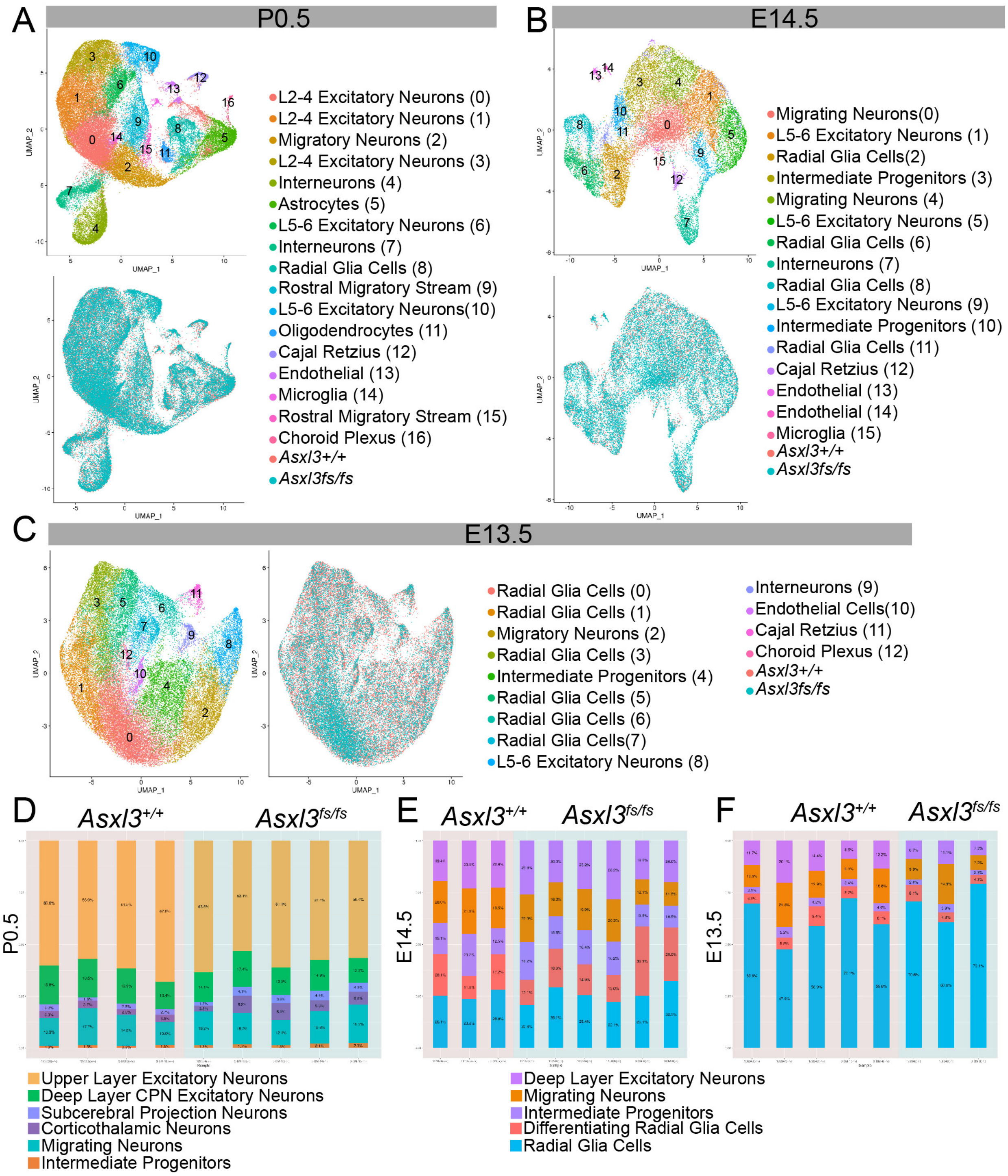
A
P0.5
L2-4 Excitatory Neurons (0)
L2-4 Excitatory Neurons (1)
Migratory Neurons (2)
L2-4 Excitatory Neurons (3)
Interneurons (4)
Astrocytes (5)
L5-6 Excitatory Neurons (6)
Interneurons (7)
Radial Glia Cells (8)
Rostral Migratory Stream (9)
L5-6 Excitatory Neurons(10)
Oligodendrocytes (11)
Cajal Retzius (12)
Endothelial (13)
Microglia (14)
Rostral Migratory Stream (15)
Choroid Plexus (16)
Asxl3+/+
Asxl3fs/fs
B
E14.5
Migrating Neurons(0)
L5-6 Excitatory Neurons (1)
Radial Glia Cells(2)
Intermediate Progenitors (3)
Migrating Neurons (4)
L5-6 Excitatory Neurons (5)
Radial Glia Cells (6)
Interneurons (7)
Radial Glia Cells (8)
L5-6 Excitatory Neurons (9)
Intermediate Progenitors (10)
Radial Glia Cells (11)
Cajal Retzius (12)
Endothelial (13)
Endothelial (14)
Microglia (15)
Asxl3+/+
Asxl3fs/fs
C
E13.5
Radial Glia Cells (0)
Radial Glia Cells (1)
Migratory Neurons (2)
Radial Glia Cells (3)
Intermediate Progenitors (4)
Radial Glia Cells (5)
Radial Glia Cells (6)
Radial Glia Cells(7)
L5-6 Excitatory Neurons (8)
Interneurons (9)
Endothelial Cells(10)
Cajal Retzius (11)
Choroid Plexus (12)
Asxl3+/+
Asxl3fs/fs
UMAP_1
UMAP_2
D
Asxl3+/+
Asxl3fs/fs
P0.5
E
Asxl3+/+
Asxl3fs/fs
E14.5
F
Asxl3+/+
Asxl3fs/fs
E13.5
Upper Layer Excitatory Neurons
Deep Layer CPN Excitatory Neurons
Subcerebral Projection Neurons
Corticothalamic Neurons
Migrating Neurons
Intermediate Progenitors
Deep Layer Excitatory Neurons
Migrating Neurons
Intermediate Progenitors
Differentiating Radial Glia Cells
Radial Glia Cells

**Supplemental Figure 4. Cortical cell types profiled with Seq-Well A,** Unsupervised clustering of P0.5 cells collected from *Asxl3*$^{+/+}$ (23,511 cells; *n*=4) and *Asx3*$^{fs/fs}$ (44,384 cells; *n*=6) samples after removal of blood cells. Top UMAP color coded by cell type and bottom UMAP colored by genotype with *Asxl3*$^{+/+}$ colored red and *Asxl3*$^{fs/fs}$ colored blue. Cell type annotations are located to the right. **B,** Unsupervised clustering of E14.5 cells collected from *Asxl3*$^{+/+}$ (6,669 cells; *n*=5) and *Asx3*$^{fs/fs}$ (18,294 cells; *n*=7) samples after removal of blood cells. Top UMAP color coded by cell type and bottom UMAP colored by genotype with *Asxl3*$^{+/+}$ colored red and *Asxl3*$^{fs/fs}$ colored blue. Cell type annotations are located to the right. **C,** Unsupervised clustering of E13.5 cells collected from *Asxl3*$^{+/+}$ (26,780 cells; *n*=5) and *Asx3*$^{fs/fs}$ (20,140 cells; *n*=3) samples after removal of blood cells. Left UMAP color coded by cell type and right UMAP colored by genotype with *Asxl3*$^{+/+}$ colored red and *Asxl3*$^{fs/fs}$ colored blue. Cell type annotations are located to the right. Stacked barplots depicting distribution of detected excitatory cell types within each sample at **D,** P0.5 (left), **E,** E14.5 (center), **F,** E13.5 (right). Barplots are organized by genotype with *Asxl3*$^{+/+}$ samples on a red background and *Asxl3*$^{fs/fs}$ on a blue background.

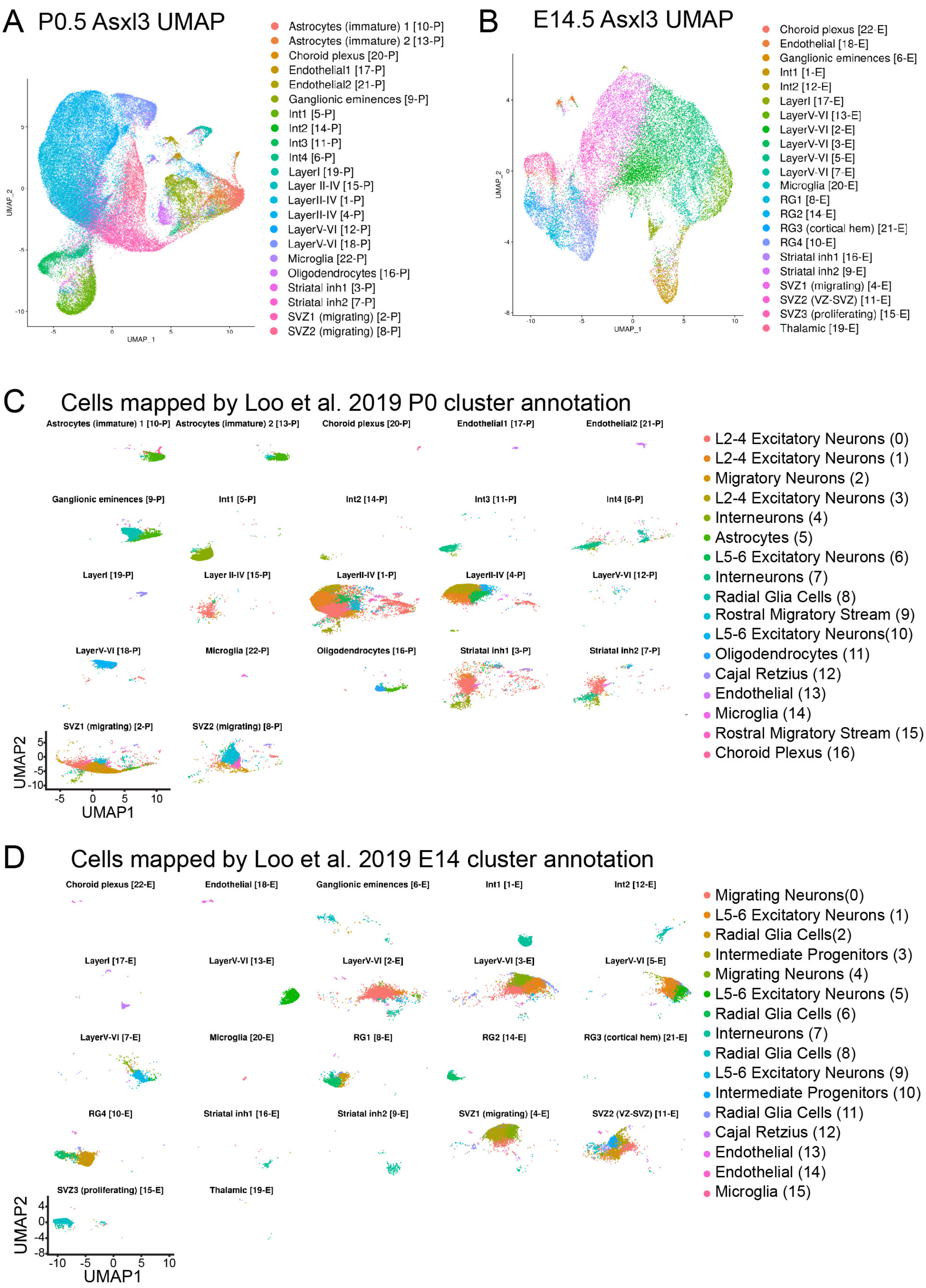
A P0.5 Asxl3 UMAP
UMAP_2
UMAP_1
Astrocytes (immature) 1 [10-P]
Astrocytes (immature) 2 [13-P]
Choroid plexus [20-P]
Endothelial1 [17-P]
Endothelial2 [21-P]
Ganglionic eminences [9-P]
Int1 [5-P]
Int2 [14-P]
Int3 [11-P]
Int4 [6-P]
LayerI [19-P]
Layer II-IV [15-P]
LayerII-IV [1-P]
LayerII-IV [4-P]
LayerV-VI [12-P]
LayerV-VI [18-P]
Microglia [22-P]
Oligodendrocytes [16-P]
Striatal inh1 [3-P]
Striatal inh2 [7-P]
SVZ1 (migrating) [2-P]
SVZ2 (migrating) [8-P]
B E14.5 Asxl3 UMAP
UMAP_2
UMAP_1
Choroid plexus [22-E]
Endothelial [18-E]
Ganglionic eminences [6-E]
Int1 [1-E]
Int2 [12-E]
LayerI [17-E]
LayerV-VI [13-E]
LayerV-VI [2-E]
LayerV-VI [3-E]
LayerV-VI [5-E]
LayerV-VI [7-E]
Microglia [20-E]
RG1 [8-E]
RG2 [14-E]
RG3 (cortical hem) [21-E]
RG4 [10-E]
Striatal inh1 [16-E]
Striatal inh2 [9-E]
SVZ1 (migrating) [4-E]
SVZ2 (VZ-SVZ) [11-E]
SVZ3 (proliferating) [15-E]
Thalamic [19-E]
C Cells mapped by Loo et al. 2019 P0 cluster annotation
Astrocytes (immature) 1 [10-P]
Astrocytes (immature) 2 [13-P]
Choroid plexus [20-P]
Endothelial1 [17-P]
Endothelial2 [21-P]
Ganglionic eminences [9-P]
Int1 [5-P]
Int2 [14-P]
Int3 [11-P]
Int4 [6-P]
LayerI [19-P]
Layer II-IV [15-P]
LayerII-IV [1-P]
LayerII-IV [4-P]
LayerV-VI [12-P]
LayerV-VI [18-P]
Microglia [22-P]
Oligodendrocytes [16-P]
Striatal inh1 [3-P]
Striatal inh2 [7-P]
SVZ1 (migrating) [2-P]
SVZ2 (migrating) [8-P]
UMAP2
UMAP1
L2-4 Excitatory Neurons (0)
L2-4 Excitatory Neurons (1)
Migratory Neurons (2)
L2-4 Excitatory Neurons (3)
Interneurons (4)
Astrocytes (5)
L5-6 Excitatory Neurons (6)
Interneurons (7)
Radial Glia Cells (8)
Rostral Migratory Stream (9)
L5-6 Excitatory Neurons(10)
Oligodendrocytes (11)
Cajal Retzius (12)
Endothelial (13)
Microglia (14)
Rostral Migratory Stream (15)
Choroid Plexus (16)
D Cells mapped by Loo et al. 2019 E14 cluster annotation
Choroid plexus [22-E]
Endothelial [18-E]
Ganglionic eminences [6-E]
Int1 [1-E]
Int2 [12-E]
LayerI [17-E]
LayerV-VI [13-E]
LayerV-VI [2-E]
LayerV-VI [3-E]
LayerV-VI [5-E]
LayerV-VI [7-E]
Microglia [20-E]
RG1 [8-E]
RG2 [14-E]
RG3 (cortical hem) [21-E]
RG4 [10-E]
Striatal inh1 [16-E]
Striatal inh2 [9-E]
SVZ1 (migrating) [4-E]
SVZ2 (VZ-SVZ) [11-E]
SVZ3 (proliferating) [15-E]
Thalamic [19-E]
UMAP2
UMAP1
Migrating Neurons(0)
L5-6 Excitatory Neurons (1)
Radial Glia Cells(2)
Intermediate Progenitors (3)
Migrating Neurons (4)
L5-6 Excitatory Neurons (5)
Radial Glia Cells (6)
Interneurons (7)
Radial Glia Cells (8)
L5-6 Excitatory Neurons (9)
Intermediate Progenitors (10)
Radial Glia Cells (11)
Cajal Retzius (12)
Endothelial (13)
Endothelial (14)
Microglia (15)

**Supplemental Figure 5. Alignment of *Asxl3* datasets with E14.5 and P0 Loo et. al 2019 open source data** UMAP of our **A,** P0.5 and **B,** E14.5 scRNA-seq data with colors and clustering assignment derived from Loo et al. 2019 P0.5 and E14.5 annotations. **C,** P0.5 and **D,** E14.5 UMAP plots displaying cells that align to previously published cell types detected in Loo et al. 2019 P0.5 and E14.5 datasets.

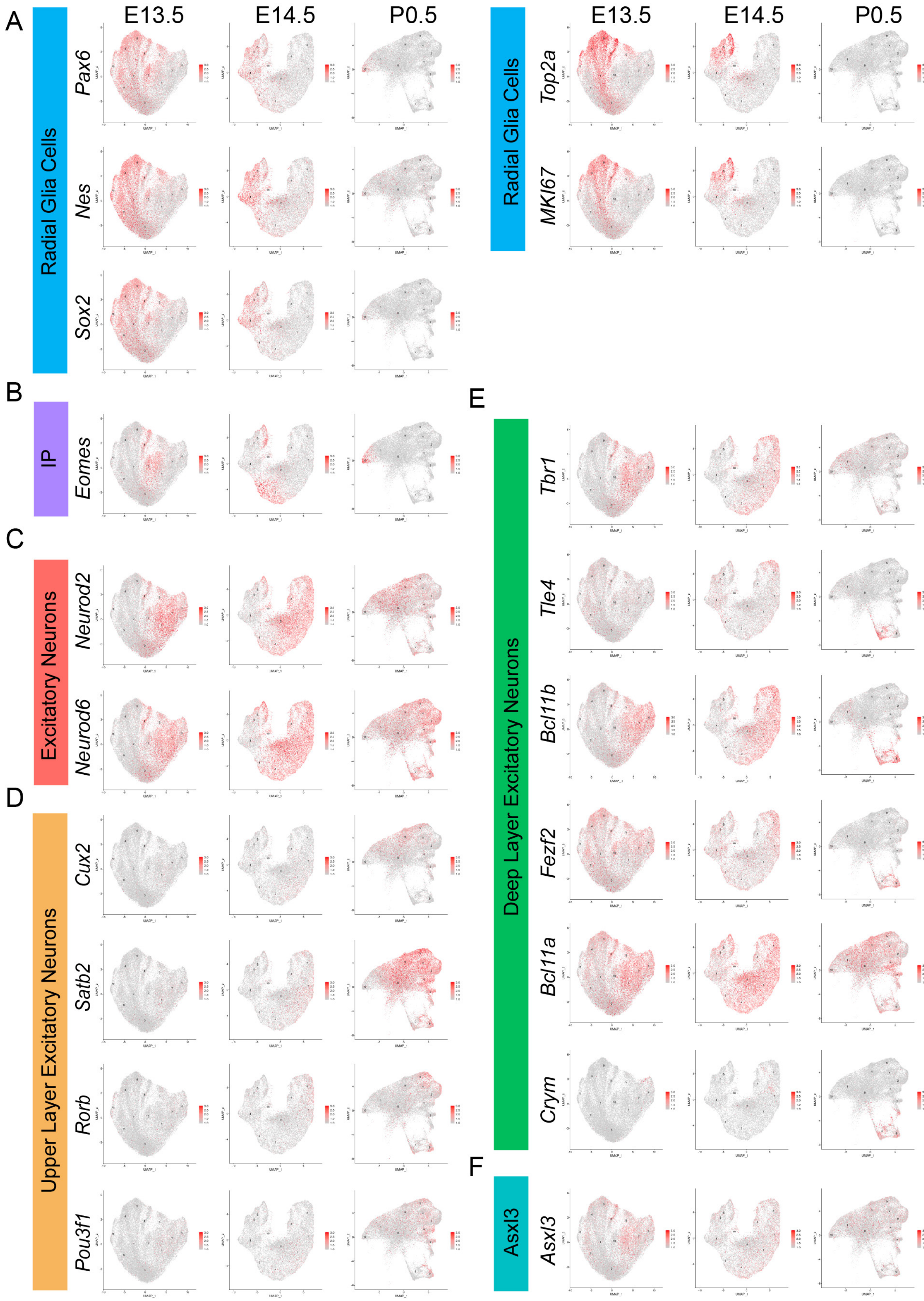

A
B
C
D
E
F
E13.5
E14.5
P0.5
Radial Glia Cells
Pax6
Nes
Sox2
Top2a
MKI67
IP
Eomes
Excitatory Neurons
Neurod2
Neurod6
Upper Layer Excitatory Neurons
Cux2
Satb2
Rorb
Pou3f1
Deep Layer Excitatory Neurons
Tbr1
Tle4
Bcl11b
Fezf2
Bcl11a
Crym
Asxl3

**Supplemental Figure 6. Characterization of E13.5, E14.5, and P0.5 excitatory cell types** E13.5, E14.5, and P0.5 feature plots showing the expression of known markers for **A,** radial glia cells, **B,** intermediate progenitors (IP), **C,** excitatory neurons, **D,** upper layer excitatory neurons, and **E,** deep layer excitatory neurons. **F,** E13.5, E14.5, and P0.5 feature plots showing the expression of *Asxl3*.

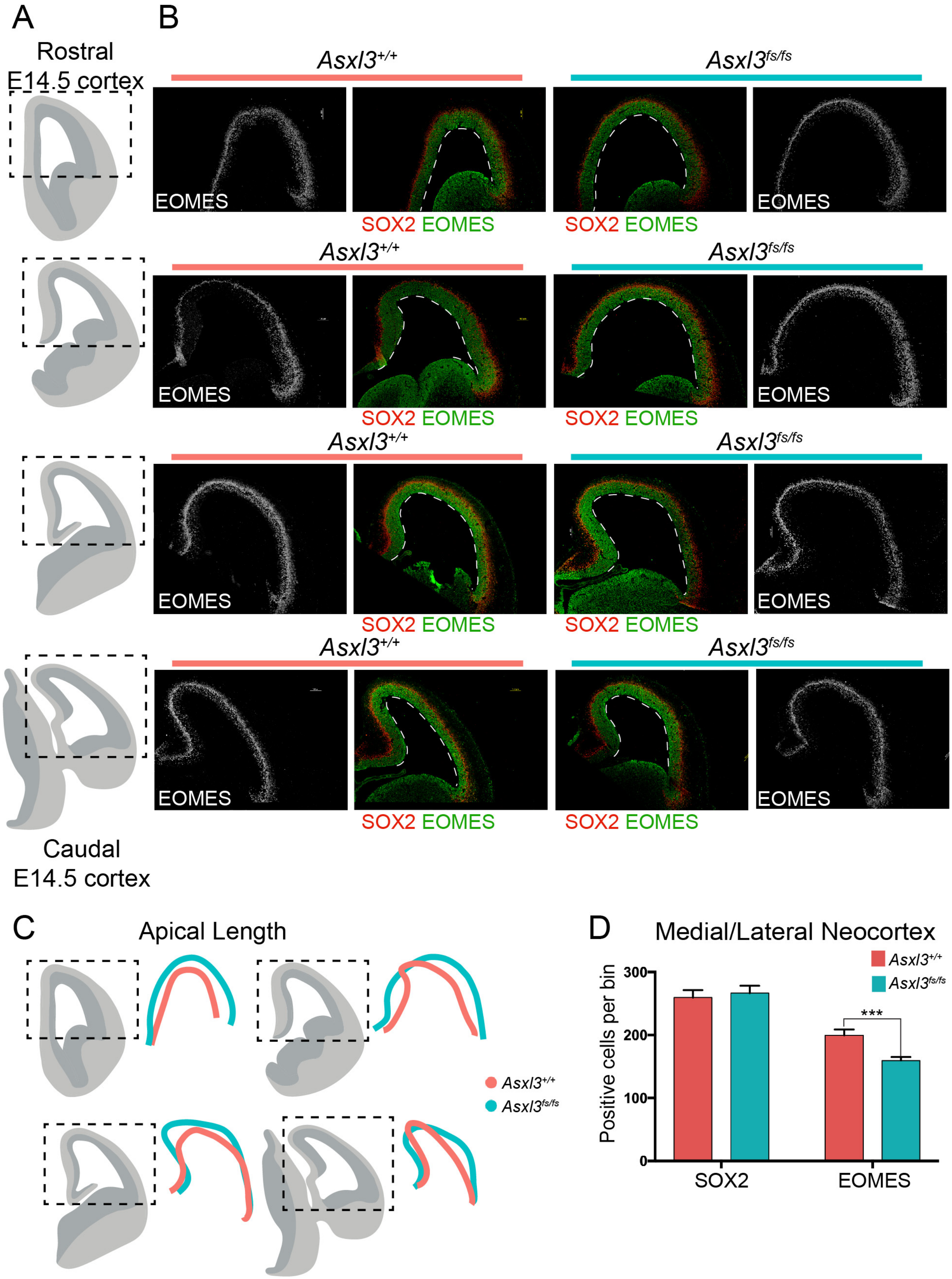
A
Rostral
E14.5 cortex
B
Asxl3+/+
Asxl3fs/fs
EOMES
SOX2 EOMES
Caudal
E14.5 cortex
C
Apical Length
Asxl3+/+
Asxl3fs/fs
D
Medial/Lateral Neocortex
Positive cells per bin
300
200
100
0
***
SOX2
EOMES

**Supplemental Figure 7. Expansion of NPCs A,** Schematic illustrating coronal sections of rostral to caudal neocortical regions used for immunohistochemistry and apical length quantification. **B,** Immunohistochemical staining of *Asxl3*$^{+/+}$ (left) and *Asx3*$^{fs/fs}$ (right) E14.5 coronal cortical sections with SOX2 (Green) and EOMES (grey, red) at different regions of the rostrocaudal axis. **C,** Comparison of *Asxl3*$^{+/+}$ (red) and *Asxl3*$^{fs/fs}$ (blue) apical lengths across the E14.5 neocortex. **D,** Quantification of the number of cells expressing SOX2 or EOMES in the medial/lateral neocortex in *Asxl3*$^{+/+}$ ($n$=6) and *Asx3*$^{fs/fs}$ ($n$=6) mice. $p$=0.699 (SOX2), $p$=1.8x10$^{-4}$ (EOMES) using two-tailed unpaired Student's $t$ test. *$p$<0.05, **$p$<0.01, ***$p$<0.001.

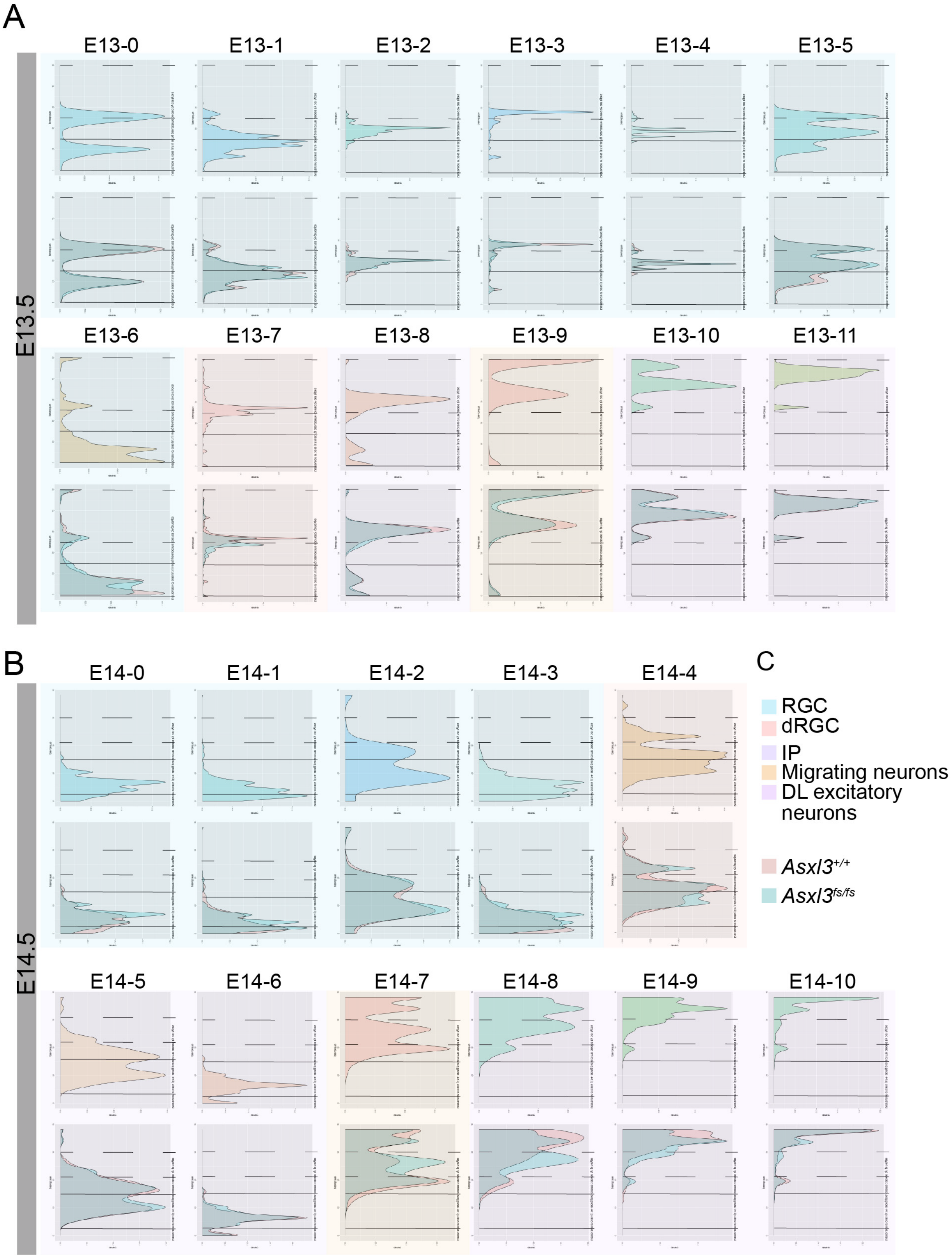
A
E13.5
E13-0
E13-1
E13-2
E13-3
E13-4
E13-5
E13-6
E13-7
E13-8
E13-9
E13-10
E13-11
B
E14.5
E14-0
E14-1
E14-2
E14-3
E14-4
E14-5
E14-6
E14-7
E14-8
E14-9
E14-10
C
RGC
dRGC
IP
Migrating neurons
DL excitatory neurons
Asxl3+/+
Asxl3fs/fs

**Supplemental Figure 8. E13.5 and E14.5 pseudotime analysis by cluster** Monocle 3 pseudotime analysis showing the pseudotime histograms for individual **A,** E13.5 and **B,** E14.5 clusters. The top graph shows the psuedotime ordering for the entire cluster. The bottom graph is colored by genotype (Asxl3$^{+/+}$, red; Asxl3$^{fs/fs}$, blue). Pseudotime maps are grouped by cell types. **C,** Legend depicting the cell type colors.

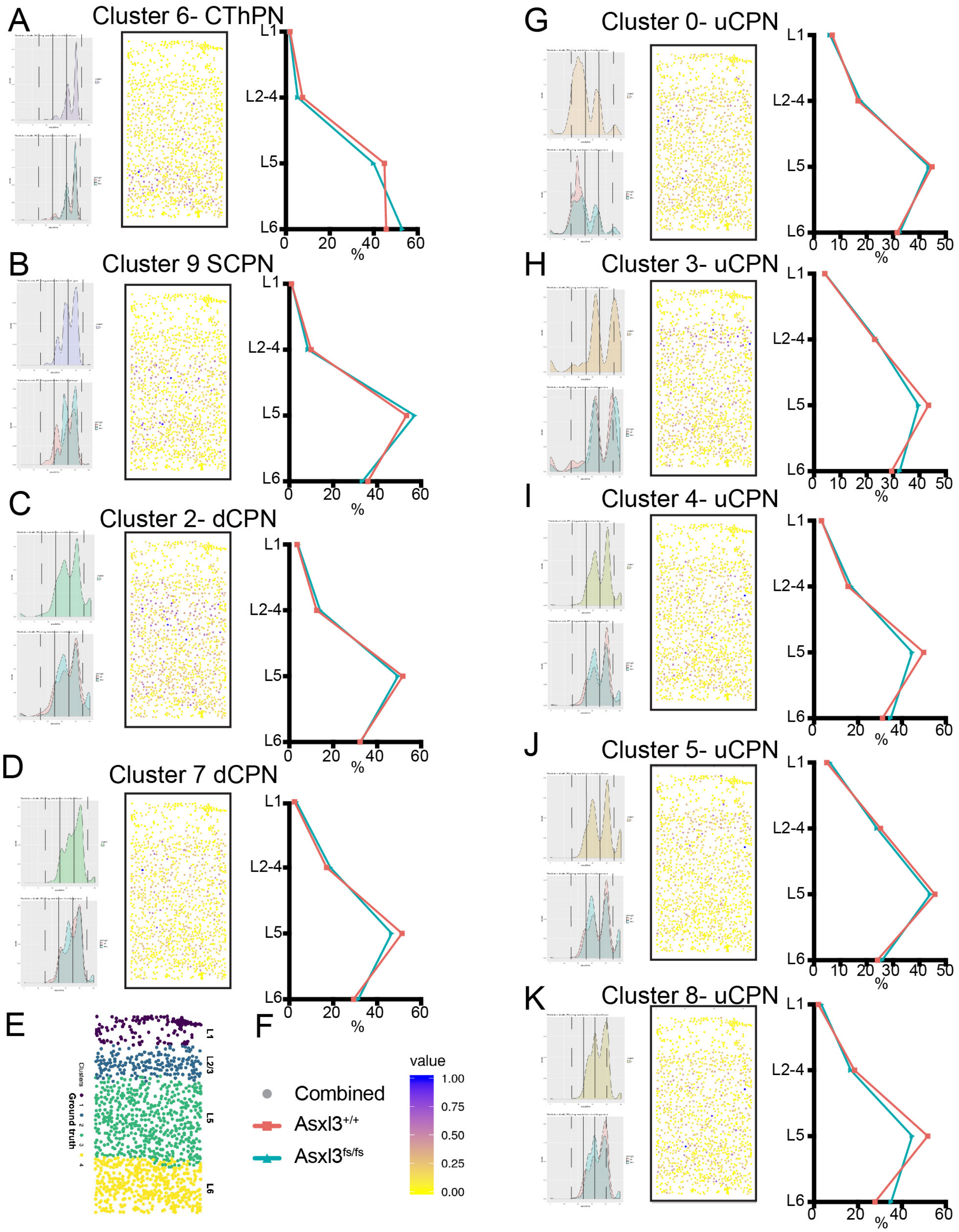
A
Cluster 6- CThPN
B
Cluster 9 SCPN
C
Cluster 2- dCPN
D
Cluster 7 dCPN
E
F
G
Cluster 0- uCPN
H
Cluster 3- uCPN
I
Cluster 4- uCPN
J
Cluster 5- uCPN
K
Cluster 8- uCPN
L1
L2-4
L5
L6
L2/3
%
Clusters
Ground truth
Combined
Asxl3+/+
Asxl3fs/fs
value
1.00
0.75
0.50
0.25
0.00

**Supplemental Figure 9. Tangram mapping of P0.5 *Asxl3*$^{fs/fs}$ scRNA-seq data to STARmap spatial transcriptomic data** Left: Monocle 3 pseudotime analysis showing the pseudotime histograms for individual P0.5 clusters identified by scRNA-seq. Middle: probabilistic mapping of our single cell gene expression data onto STARmap spatial gene expression data (Wang et al. 2018) using Tangram (Biancalani et al. 2020). Dots colored by the probability of our cell type mapping to the STARmap cell type (blue high probability, yellow low probability). Right: Distribution of cells mapping to spatial gene expression data for L6, L5, L2-4, and L1. Each analysis was performed for **A,** cluster P0-6 CThPN cells **B,** cluster P0-9 SCPN, **C,** cluster P0-2 dCPN, **D,** cluster P0-7 dCPN, **G,** Cluster P0-0 uCPN, **H,** Cluster P0-3 uCPN, **I,** Cluster P0-4 uCPN, **J,** Cluster P0-5 uCPN, **K,** Cluster P0-8 uCPN. Distributions are colored by genotype (*Asxl3*$^{+/+}$ red, *Asxl3*$^{fs/fs}$ blue). **E,** Spatial localization of cell types. **F,** Figure legend for corresponding plots and graphs.
